# Bispecific VSV-G Engagers: A Platform for Cell-Type-Specific Delivery of Viral Vectors

**DOI:** 10.64898/2026.11.02.763599

**Authors:** Gopal Naik Nenavath, Diana Gataulin, Miriam Eisenstein, Gideon Schreiber, Md Sharif Hasan, Nandakumar Packiriswamy, Pragati Jain, Tatenda Kadungure, Harshitha Anantharama, Chia-Hsuan Chin, Aravind Kancharla, Hamid Salimi, Patrycja Lech, Luis Blancas-Mejia, Miguel Angel Muñoz-Alía, Kah-Whye Peng, Stephen J. Russell, Menachem Rubinstein

## Abstract

VSV-G-pseudotyped lentiviral vectors (LVVs) enable highly efficient ex vivo gene delivery through the broadly expressed LDL receptor (LDLR) but lack the cell-type specificity required for in vivo applications. To provide cell-type specificity, we developed reversible engagers consisting of a cell-specific ligand or antibody fused to the ∼40-amino-acid VSV-G-binding domains of LDLR. Following endocytosis, the engager dissociates, releasing unmodified VSV-G for efficient delivery of the genetic cargo. An engager comprising the CEACAM8 N-terminal domain enables selective transduction of CEACAM6-positive epithelial cells. A trimeric CEACAM8-based engager matching the VSV-G homotrimeric spike blocked non-specific VSV infection through LDLR with an IC_50_ of 16.7 pM, while enabling effective VSV-mediated lysis of CEACAM6-positive tumor cells. An anti-CD3-based trimeric engager selectively transduced non-preactivated T cells in human PBMCs with high transduction efficiency. Engager-mediated in vivo CAR T cell generation resulted in sustained control of human OPM2 multiple myeloma tumors in NSG mice. This platform may further advance in vivo gene therapy, including in vivo CAR T-cell therapy and tumor-directed oncolytic virotherapy.

---

Vesicular stomatitis virus (VSV) is an enveloped virus whose negative-sense single-stranded RNA genome encodes five proteins: nucleoprotein, phosphoprotein, matrix protein, glycoprotein, and viral polymerase. The glycoprotein (VSV-G) forms trimeric transmembrane spikes that dictate receptor recognition, endocytosis, and fusion of the viral envelope with the endosomal membrane, thereby delivering its RNA and proteins into the host cell cytoplasm^1^. VSV exhibits broad tropism for many vertebrate species and most cell types, including quiescent cells^1, 2^, which has prompted the extensive use of VSV-G-pseudotyped lentiviral vectors (LVVs) for transducing various cell types. VSV-G pseudotyping also provides high transduction efficiency, enhanced vector titer, and storage stability^3^. Hematopoietic stem cells (HSCs) transduced with LVVs encoding functional genes have already been approved for the gene therapy of several hereditary diseases^4^. Similarly, T cells transduced ex vivo with LVVs encoding chimeric antigen receptors (CAR T) are in clinical use to treat B-cell leukemia and multiple myeloma. They are in clinical development for additional indications, including solid tumors^5–7^, and autoimmune diseases^8, 9^.

Despite their high efficacy, CAR T cell therapy and HSC-directed gene therapy require complex, multi-step procedures, including harvesting cells from the patient, isolating highly purified, specific cell types (e.g., HSCs or T cells), and activating the cells before LVV-mediated transduction^10^. The resulting cells are then amplified ex vivo, purified, and transfused into the same donor. Such protocols must be repeated for each patient individually to prevent immune rejection, making these therapies expensive and inaccessible to most patients. Administering systemic viral vectors can simplify gene and CAR T cell therapies, offering off-the-shelf solutions that eliminate all ex vivo steps and significantly reduce time and costs. However, the broad tropism of VSV-G-pseudotyped vectors renders them unsuitable for systemic in vivo administration. This limitation prompted the development of reengineered vectors to target specific cell types^11,12,13,14^, albeit at the cost of reduced transduction efficiency and the need to test them and obtain regulatory approval for the modified vectors.

In addition to using VSV-G-pseudotyped LVV for gene therapy and CAR T cell therapy, both natural and recombinant forms of VSV have been studied and adapted for oncolytic virotherapy^15–17^. VSV directly eliminates tumor cells and stimulates an anti-tumor immune response, providing a complementary therapeutic mechanism^18^. Many tumor cell types are deficient in their interferon response, thereby providing oncolytic virus tumor selectivity. However, following systemic VSV administration, most viral particles are lost by uptake into virus-resistant healthy cells. Selective binding of an oncolytic virus to tumor cells would enhance the safety and efficacy of oncolytic virotherapy. Recently, it was demonstrated that oncolytic VSV combined with CAR-T cell therapy was effective in treating a melanoma model in mice^19^.

VSV and VSV-G-pseudotyped vectors bind to cells and are endocytosed via the low-density lipoprotein receptor (LDLR) and its family members^20^. These receptors are evolutionarily conserved and broadly expressed, explaining the broad tropism of VSV and LVV. We have identified a soluble fragment of human LDLR (sLDLR) in cell culture supernatants, consisting of the first four N-terminal Class A repeats, which potently inhibits VSV^21^. We reported that VSV binds to the CR3 domain of sLDLR, as determined using a CR3-directed monoclonal antibody^20^. Later, this observation was confirmed and extended to CR2^22^. All CR repeats are calcium-binding proteins, and we have demonstrated that VSV-G binding to LDLR is Ca^2+^-dependent, suggesting that low endosomal Ca^2+^ levels trigger vector release, enabling fusion with the endosomal membrane^20^.

Here, we propose an approach for directing VSV- and VSV-G-pseudotyped viral vectors to specific cell types. To this end, we designed reversible bispecific engager proteins consisting of the VSV-G binding domains of LDLR fused to ligands or antibodies targeting cell-type-specific receptors (Fig. 1). This approach preserves the potency of unmodified VSV and VSV-G pseudotyped viral vectors while directing them to specific cell types.

**Fig. 1.**
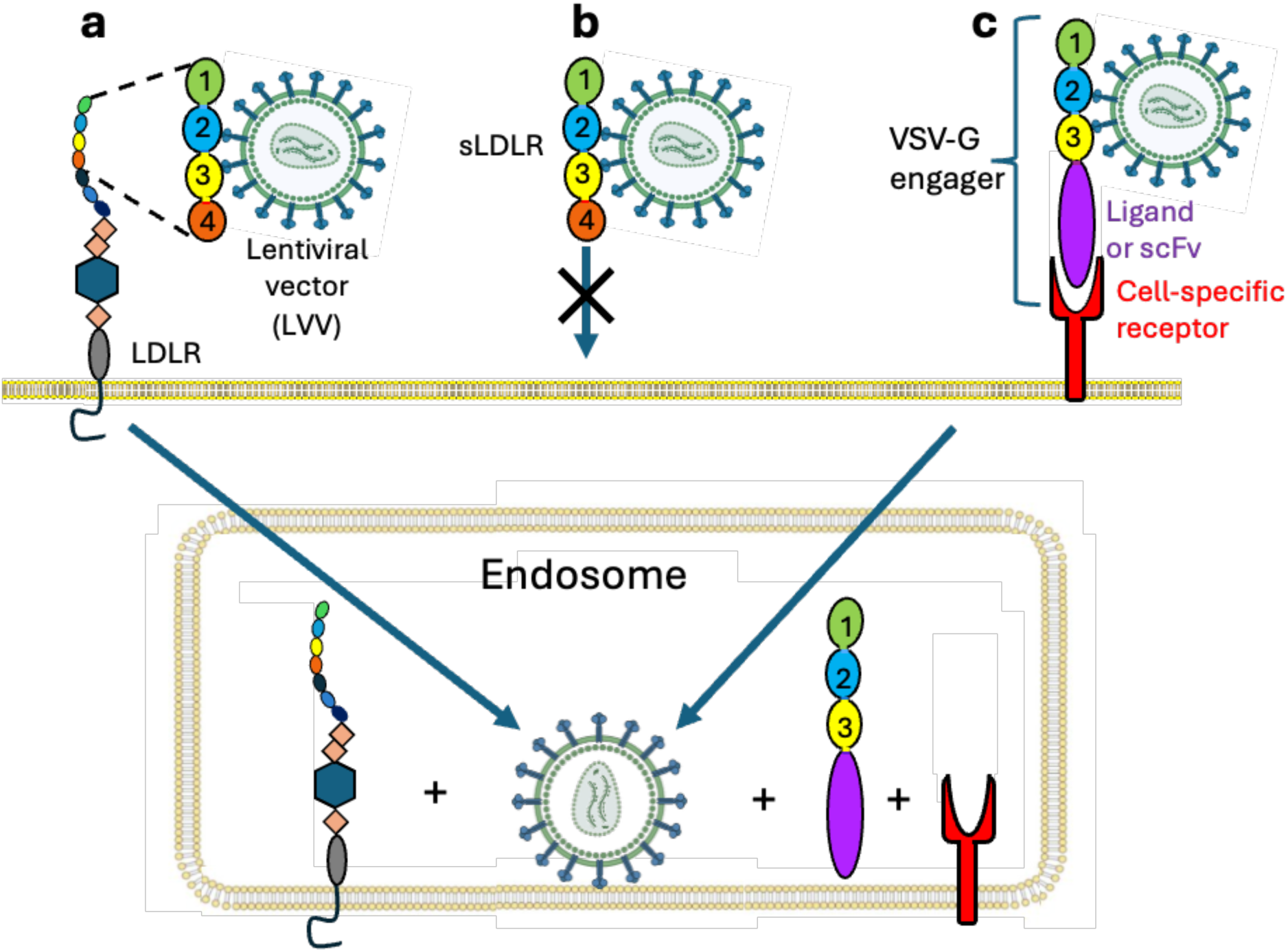
LDLR-based engagers retarget VSV and LVV. **a,** VSV and VSV-G-pseudotyped LVV bind reversibly to the N-terminal cysteine-rich (CR) repeats 2 and 3 of the broadly expressed LDLR. After endocytosis, VSV-G dissociates from LDLR and mediates fusion with the endosomal membrane. **b,** Soluble LDLR (sLDLR) inhibits VSV infection and LVV transduction by competing with cell-surface LDLR family members. **c,** A reversible engager comprises VSV-G-binding CR domains fused to a ligand or antibody for a cell-type-specific receptor. The engager directs the vector to the target receptor while blocking nonspecific entry via the LDLR. Following endocytosis, unmodified VSV or LVV is released from the engager and delivers its cargo to the cytoplasm.

## Results

### CEACAM8 engagers redirect VSV to CEACAM6-expressing tumor cells

Carcinoembryonic antigen-related cell adhesion molecule 6 (CEACAM6) is highly expressed in several epithelial cancers, including colon, ovarian, breast, and pancreatic cancers, making it a potential target for tumor-selective treatments^23, 24^. CEACAM8 and CEACAM6 form heterodimers by binding through their N-terminal domains^25, 26^. In addition, CEACAM6 serves as an entry portal for influenza viruses and certain bacteria^27, 28^, suggesting that it allows entry of large, virus-sized particles. We designed an engager consisting of sLDLR (amino acids 25-145 of human pro-LDLR) fused via a short linker to the N-terminal domain of CEACAM8 (amino acids 35-140 of human pro-CEACAM8). CEACAM8-sLDLR, an engager whose protein subunits are fused in a reversed order, was also produced and used interchangeably. (Supplementary Fig. 1a, b). These bispecific engagers and standalone sLDLR_(25-145)_ were effectively produced in HEK293T and CHO-S cells. His_6_-containing engagers were purified using Ni^2+^-NTA capture and, in some cases, by additional size-exclusion chromatography.

We evaluated the capacity of the engagers to direct VSV infection to CEACAM6-positive pancreatic adenocarcinoma AsPC-1 cells. VSV was pre-incubated with either sLDLR_(25-145)_ or CEACAM8-sLDLR before being added to these cells, and residual cell viability was assessed by Crystal violet staining at 24 hours. Standalone sLDLR effectively blocked the VSV-mediated cytolysis of both CEACAM6-positive AsPC-1 cells (Fig. 2a, c) and CEACAM6-negative FS11 fibroblasts (Fig. 2b, d). In contrast, the CEACAM8-sLDLR engager enabled cytolysis of AsPC-1 cells (Fig. 2a), while protecting the CEACAM6-negative FS11 fibroblasts^29^ (Fig. 2b).

**Fig. 2.**
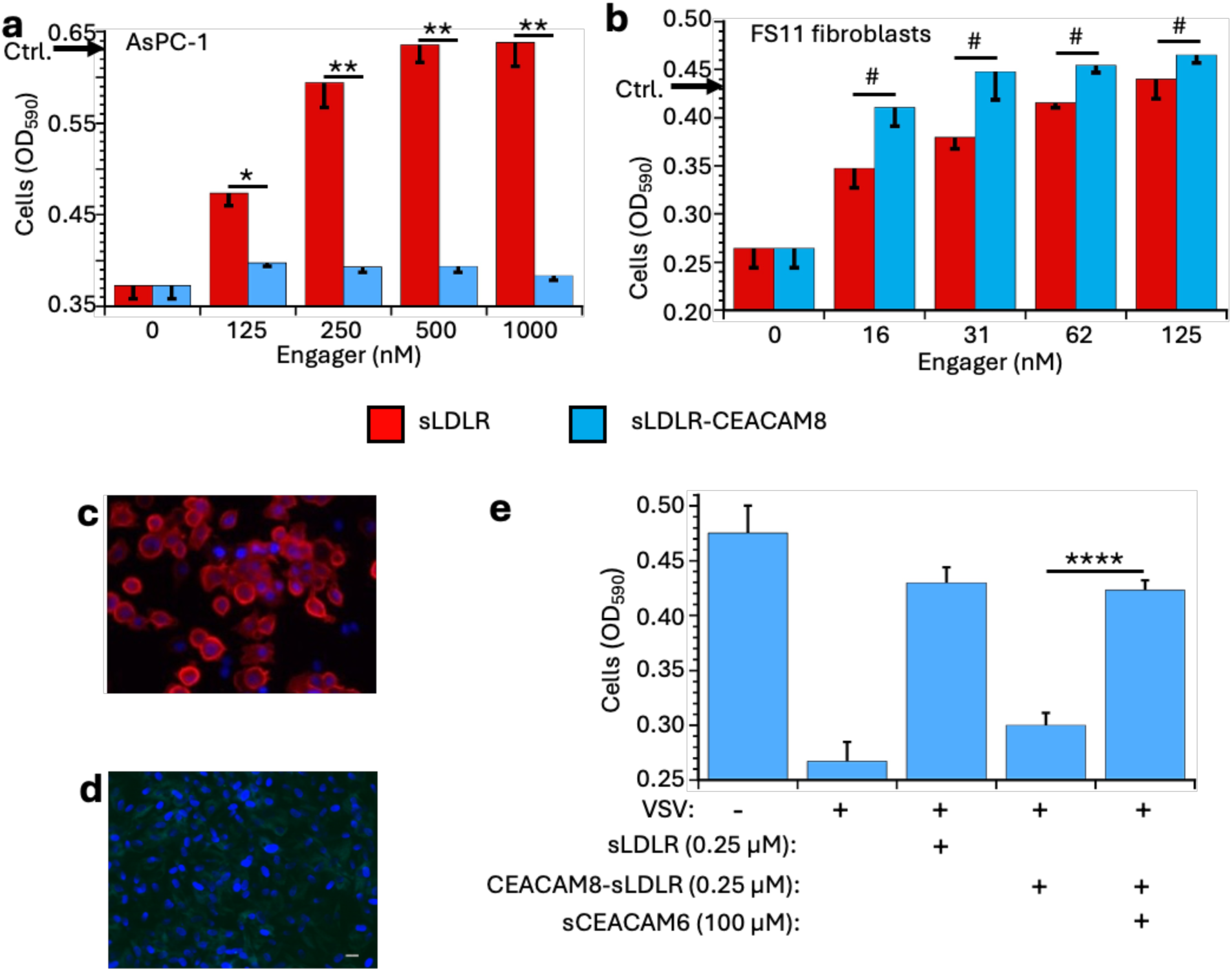
CEACAM8-based engagers redirect VSV through CEACAM6. **a,** Effect of sLDLR and CEACAM8-sLDLR on VSV-mediated cytolysis of CEACAM6-positive AsPC-1 pancreatic adenocarcinoma cells. VSV was preincubated with the indicated protein, and cell viability was measured after 24 h by crystal violet staining. Engager concentrations are the final concentrations in culture. Arrows on the Y axis indicate untreated control cultures (n = 3; *P < 0.05, **P < 0.005). **b,** Effect of sLDLR and CEACAM8-sLDLR on the viability of VSV-infected CEACAM6-negative FS11 fibroblasts under the conditions in **a** (n = 3; ^#^P > 0.12, not significant). **c, d,** Immunostaining of cell-surface CEACAM6 in AsPC-1 cells (**c**) and primary FS11 fibroblasts (**d**). **e,** Viability of AsPC-1 cells after infection with VSV preincubated with the indicated proteins, including soluble CEACAM6 competitor (n ≥ 4; ****P < 0.0001).

To further examine the pathway of vector entry in the presence of the bispecific engager, we performed a competition study using a soluble form of CEACAM6 corresponding to its N-terminal domain (amino acids 35-140 of human pro-CEACAM6). sCEACAM6 competed with cell-surface CEACAM6 for binding to the CEACAM8 domain of the engager. Mixing this competitor with the vector and engager inhibited VSV infection of AsPC-1 cells, indicating that vector entry was mediated by binding to cell-surface CEACAM6 and subsequent VSV endocytosis (Fig. 2e).

The CEACAM8-sLDLR-mediated inhibition of VSV binding to CEACAM6-negative cells exhibited a half-maximal inhibitory concentration (IC_50_) of about 3 nM. To increase the avidity of the engager for VSV-G, we designed a trimeric engager that concomitantly captures the three binding domains of the homo-trimeric VSV-G spike. This engager had the general formula: CEACAM8_(35-140)_-L1-CR2-L2-CR2-L2-CR2-His_6_ (In short: CEACAM8-(CR2)_3_). L1 is a 10-amino-acid residues linker, and L2 is 34-amino-acid residues long, spanning the distances between adjacent LDLR binding regions in the trimeric VSV-G spike (Fig. 3a, b). It was produced in CHO-S cells at a high yield and purified to greater than 90% purity by Ni-NTA column chromatography followed by size-exclusion chromatography (Supplementary Fig. 2). Pre-incubation of VSV with either sLDLR or the monomeric CEACAM8-sLDLR engager inhibited infection of CEACAM6-negative FS11 fibroblasts with similar efficacy. The trimeric CEACAM8-(CR2)_3_ engager inhibited infection with an IC_50_ of 16.7 pM, a 172-fold greater potency than that of CEACAM8-sLDLR (Fig. 3c). Infection of CEACAM6-negative HeLa cells was similarly inhibited (Supplementary Fig. 3). In contrast, this engager enabled infection and cytolysis of CEACAM6-positive AsPC-1 cells, even when added at a large molar excess (Fig. 3d).

**Fig. 3.**
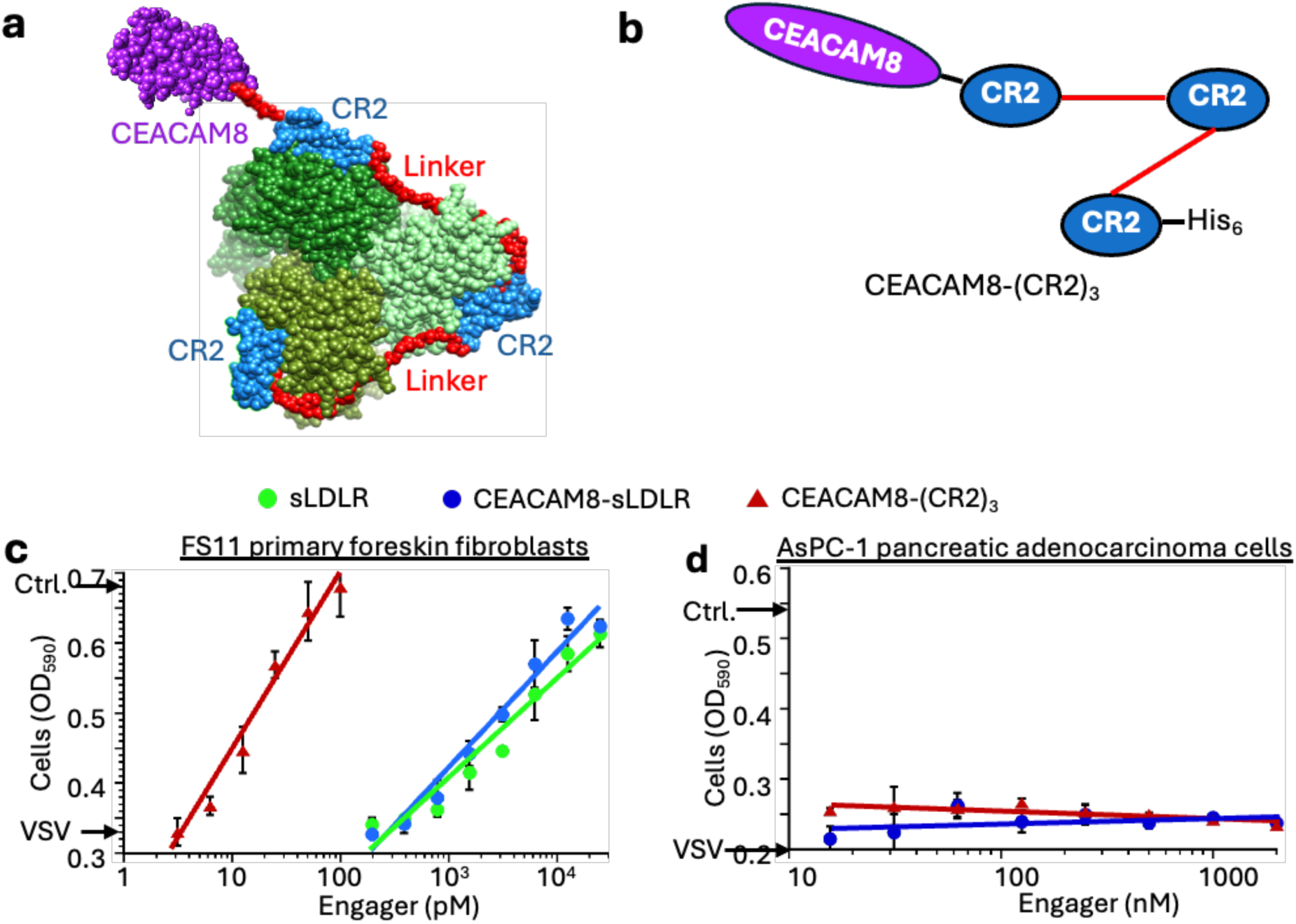
Multivalent CR2 domains increase engager potency. **a,** Space-filling model of CEACAM8-(CR2)_3_ bound to the trimeric VSV-G spike, viewed from above. CEACAM8 is shown in purple, CR domains in blue, 10- and 34-residue linkers in red, and VSV-G protomers in shades of green. **b,** Schematic of CEACAM8-(CR2)_3_. **c,** Inhibition of VSV-mediated cytolysis of CEACAM6-negative FS11 fibroblasts by sLDLR, the monomeric sLDLR-CEACAM8 and trimeric CEACAM8-(CR2)_3_ engagers. Technical triplicate dose-response curves were fitted by nonlinear regression using a four-parameter logistic model. Differences in logIC₅₀ between monomeric and trimeric engagers were evaluated by an extra-sum-of-squares F test. IC_50_ of CEACAM8-(CR2)_3_ was 16.7 pM, 95% CL = 13.9-20.2 pM. IC_50_ of CEACAM8-sLDLR was 2.9 nM, 95% CL = 2.4-3.4 nM. Potency difference = 172-fold (95% CI, 142–209). **d,** Engager-mediated VSV cytolysis of CEACAM6-positive AsPC-1 cells. VSV at MOI ∼1 was incubated with an engager for 1 h at room temperature in DMEM supplemented with 10% FBS, then added to cells in 96-well plates. Concentrations are the final ones in culture. After overnight incubation at 37 °C, cells were fixed, stained with crystal violet, and analyzed with a plate reader. Arrows on the Y-axis indicate untreated cultures and cultures exposed to VSV alone (n = 4).

### Selective engager-mediated transduction of epithelial cells

Polarized epithelial cells express CEACAM6 selectively on their apical side^30, 31^. We tested whether sLDLR-CEACAM8 would enable selective transduction of the AsPC-1 and Caco-2 epithelial-like cell lines. To this end, we pre-incubated EGFP-encoding LVV with either sLDLR or sLDLR-CEACAM8 and then transduced CEACAM6-positive AsPC-1 and Caco-2 cells with the LVV-engager mixture. The CEACAM6-negative FS11 fibroblasts served as controls. In the presence of the bispecific engager, LVV effectively transduced the CEACAM6-positive AsPC-1 and Caco-2 cells (Fig. 4a to c). In contrast, sLDLR-CEACAM8 effectively inhibited the transduction of the CEACAM6-negative FS11 fibroblasts (Fig. 4d). As previously observed^20^, sLDLR blocked transduction in all three cell types (**Fig. 4**). An engager with reversed sequences, CEACAM8-sLDLR, gave similar results (data not shown). These experiments indicated that blocking the VSV-G spike of LVV with the engager selectively transduces epithelial cells via their cell-surface CEACAM6.

**Fig. 4.**
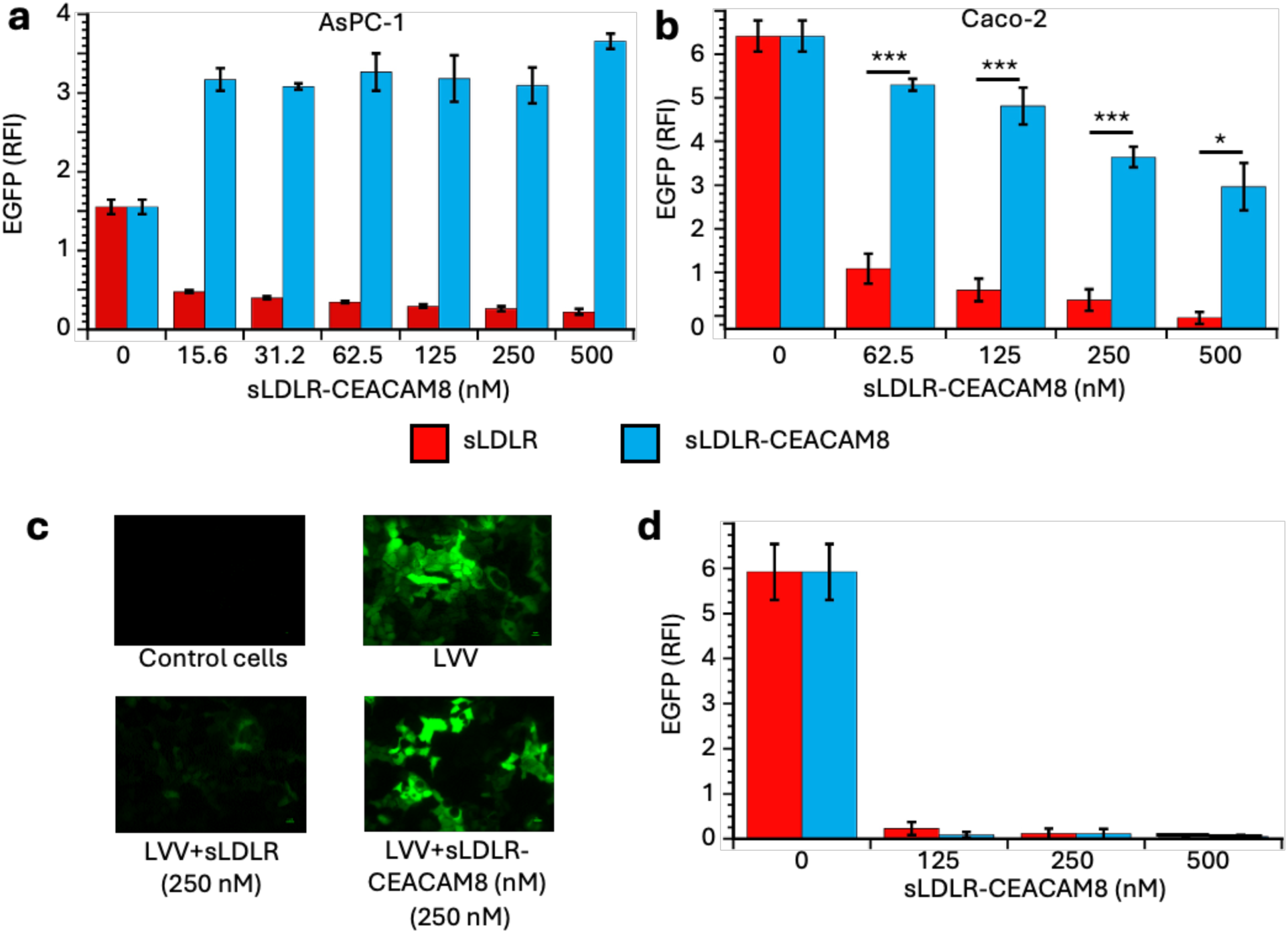
CEACAM8 engagers target LVV transduction to CEACAM6-positive cells. **a,** Relative fluorescence intensity of AsPC-1 cells transduced with EGFP-encoding LVV preincubated with sLDLR or sLDLR-CEACAM8 (n = 3; P < 0.005 at all concentrations). **b,** Relative fluorescence intensity of Caco-2 cells treated as in **a** (n = 3; *P < 0.05, ***P < 0.002). **c,** Representative fluorescence images of transduced Caco-2 cells. **d,** Relative fluorescence intensity of CEACAM6-negative FS11 fibroblasts treated as in **a** (n = 3). Differences in EGFP expression between matched sLDLR- and sLDLR-CEACAM8-treated cells were not significant.

### Design and evaluation of a CD3-directed trimeric engager

We then designed and produced a second type of trimeric engager, Engager3, incorporating sLDLR_(25-145)_ (CR1, CR2, CR3) linked to UCHT1, a CD3-targeting domain, via a trimerizing peptide (the leucine zipper domain of GCN4, *VL*). A control monomeric engager, Engager1, containing a non-trimerizing peptide of the same length, was similarly produced (Supplementary Fig. 4a). Following purification, protein quality was confirmed via SDS-PAGE and SEC-HPLC (Supplementary Fig. 4b). Unlike other approaches, this reversible engager is designed for in vivo transduction of T cells by unmodified LVV (Fig. 5).

**Fig. 5.**
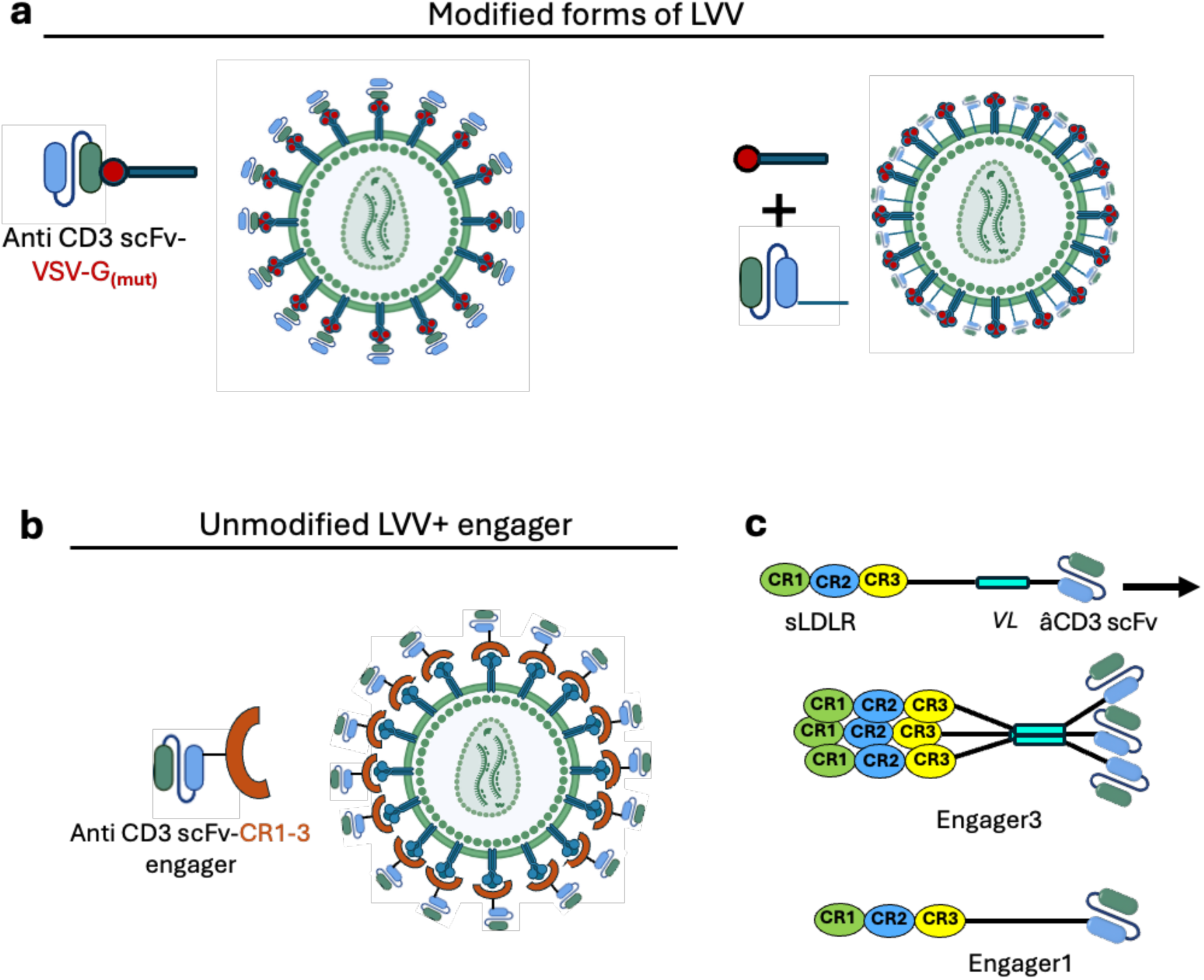
Reversible engagers enable T cell-specific transduction by unmodified VSV-G. **a,** Conventional retargeting strategies employ mutated VSV-G to reduce LDLR binding and display a CD3-directed antibody either as a VSV-G fusion or as a co-expressed envelope component. **b,** In the present approach, unmodified VSV-G-pseudotyped LVV is complexed with a reversible bispecific trimeric Engager3. **c,** Schematic representation of the trimeric sLDLR-based anti-CD3 engager used for CAR T cell generation and of the corresponding monomeric Engager1.

We then assessed the binding affinities of the adapter proteins to CD3ε/γ subunits using biolayer interferometry (BLI). The binding of monomeric Engager1 corresponded to a 1:1 binding model with a *Kd* = 40.9 ± 7.07 nM (Fig. 6a). In the absence of a trivalent binding model, a 1:1 binding model was used to calculate an apparent *Kd* = 2.2 ± 0.58 nM for Engager3, reflecting a reduced dissociation rate. To evaluate the specificity of the CD3-targeted engager, Jurkat cells and TCR-deficient Jurkat (TCR-KO) cells were transduced with pre-formed complexes of EGFP-encoding LVV with varying concentrations of trimeric or monomeric engagers. In the absence of an engager, LVV transduced both cell types. In contrast, at 10 nM, the trimeric engager selectively transduced the TCR-positive Jurkat cells with no transduction of the TCR-KO cells, suggesting the specificity of the trimeric-engager-bound LVV (Fig. 6b). The trimeric engager was 25 times more potent than the monomeric one as an inhibitor of LDLR-mediated transduction, as evaluated in the TCR-KO cells (Fig. 6c). Importantly, the overall transduction efficiency in the TCR^+^ Jurkat cells was unaffected by an excess of either engager (Fig. 6d). Furthermore, at 6-10 nM, the trimeric engager inhibited transduction of LDLR-expressing B-ALL NALM6 cells (Fig. 6e). Taken together, these results indicate that the CD3-targeted trimeric Engager3 efficiently binds to VSV-G spikes on LVV, effectively masking them from native LDLR-directed transduction, and redirecting the vector specifically to CD3-positive cells.

**Fig. 6.**
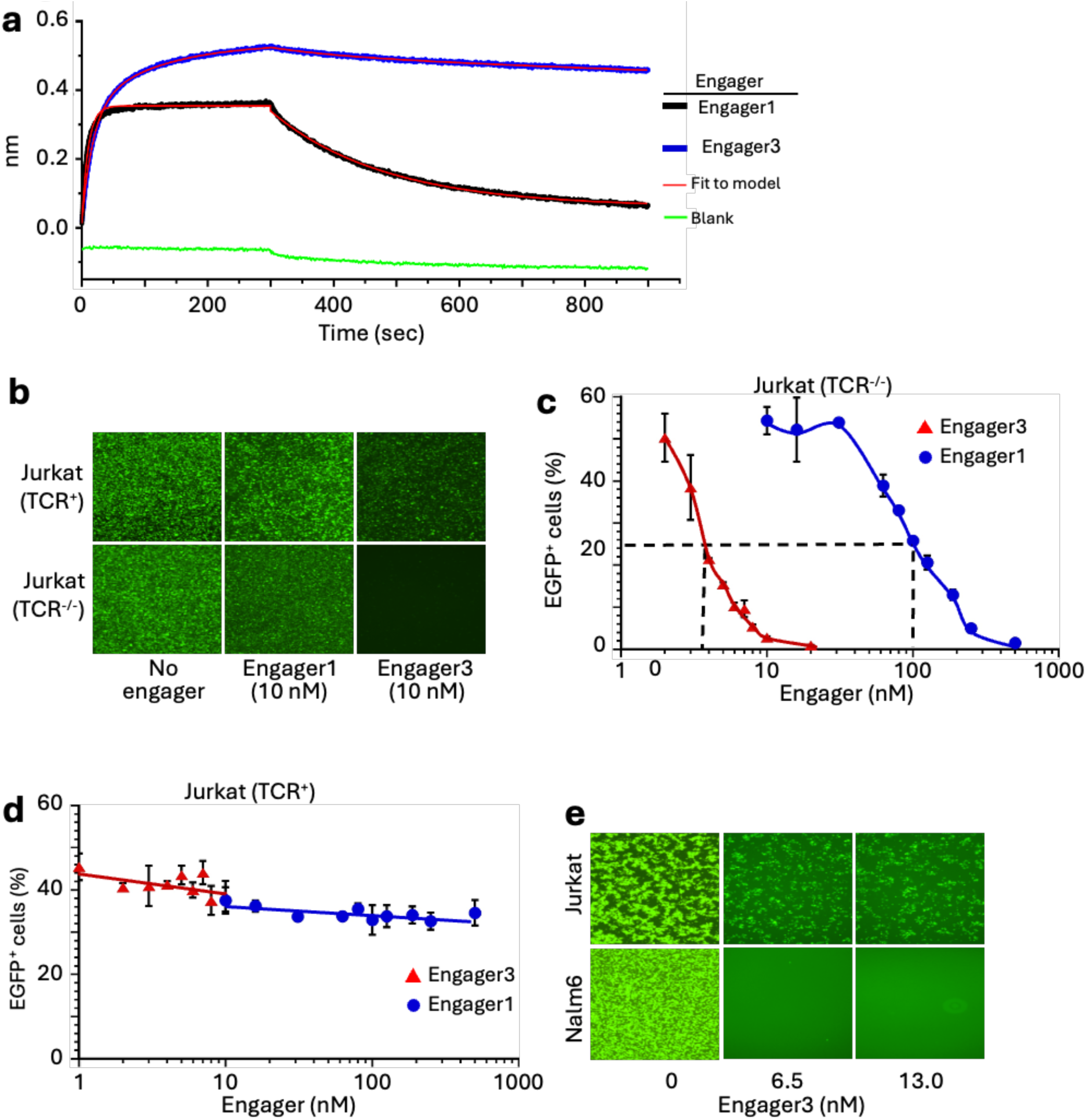
The trimeric engager mediates CD3-selective transduction. **a,** Binding of monomeric Engager1 and trimeric Engager3s to a recombinant human CD3ε/CD3γ measured by biolayer interferometry. The binding data are averages of 20 readings. A binding-model fit is shown in red. The binding of monomeric engager fits a 1:1 binding model, whereas the binding of the trimeric engager best fits a 2:1 (analyte:ligand) binding model due to the lack of a trivalent binding model. **b,** Representative fluorescence images of cells transduced with EGFP-encoding LVV complexed with 10 nM engager. **c, d,** Transduction of TCR-knockout (**c**) and TCR-positive (**d**) Jurkat cells with EGFP-encoding LVV preincubated with the indicated engager concentrations. EGFP-positive cells were quantified by flow cytometry (n = 3). **e,** Representative images of TCR-positive Jurkat cells and CD3-negative NALM6 B-acute lymphoblastic leukemia cells exposed to EGFP-encoding LVV alone or pre-incubated with the trimeric anti-CD3 engager. For all calculations, the molarity of the trimeric engager is based on its monomeric subunit.

### Highly efficient and specific engager-mediated generation of BCMA-CAR T cells in PBMC

To evaluate T-cell-specific targeting and BCMA-CAR engineering, Primary human peripheral blood mononuclear cells (PBMCs) were transduced with BCMA-CAR-encoding LVV premixed with several concentrations of the trimeric Engager3. Flow cytometry of CD3⁺, CD45⁺ cells seven days post-transduction revealed robust BCMA-CAR expression following treatment with engager-coated LVV (Fig. 7a, b). BCMA-CAR expression increased dose-dependently with the engager protein concentration, peaking at intermediate levels of 6.7-13.7 nM. At these optimal concentrations, CAR positivity reached 60-90% of the CD3^+^ cells. At concentrations exceeding ∼50 nM, CAR expression declined, likely due to competitive inhibition by excess free engager (Fig. 7c). Both CD4^+^ and CD8^+^ T cells were transduced (Fig. 7d), and T cells exhibited high CD25 expression (55–80%) at intermediate concentrations of about 7–13 nM (Fig. 7e). This indicates a sustained activation state following vector exposure. Collectively, these data demonstrate that LVV complexed with the trimeric Engager3 effectively generates BCMA CAR T cells and promotes T-cell activation. Optimal vector performance and subset modulation were consistently observed at intermediate engager protein concentrations of 6–13 nM.

**Fig. 7.**
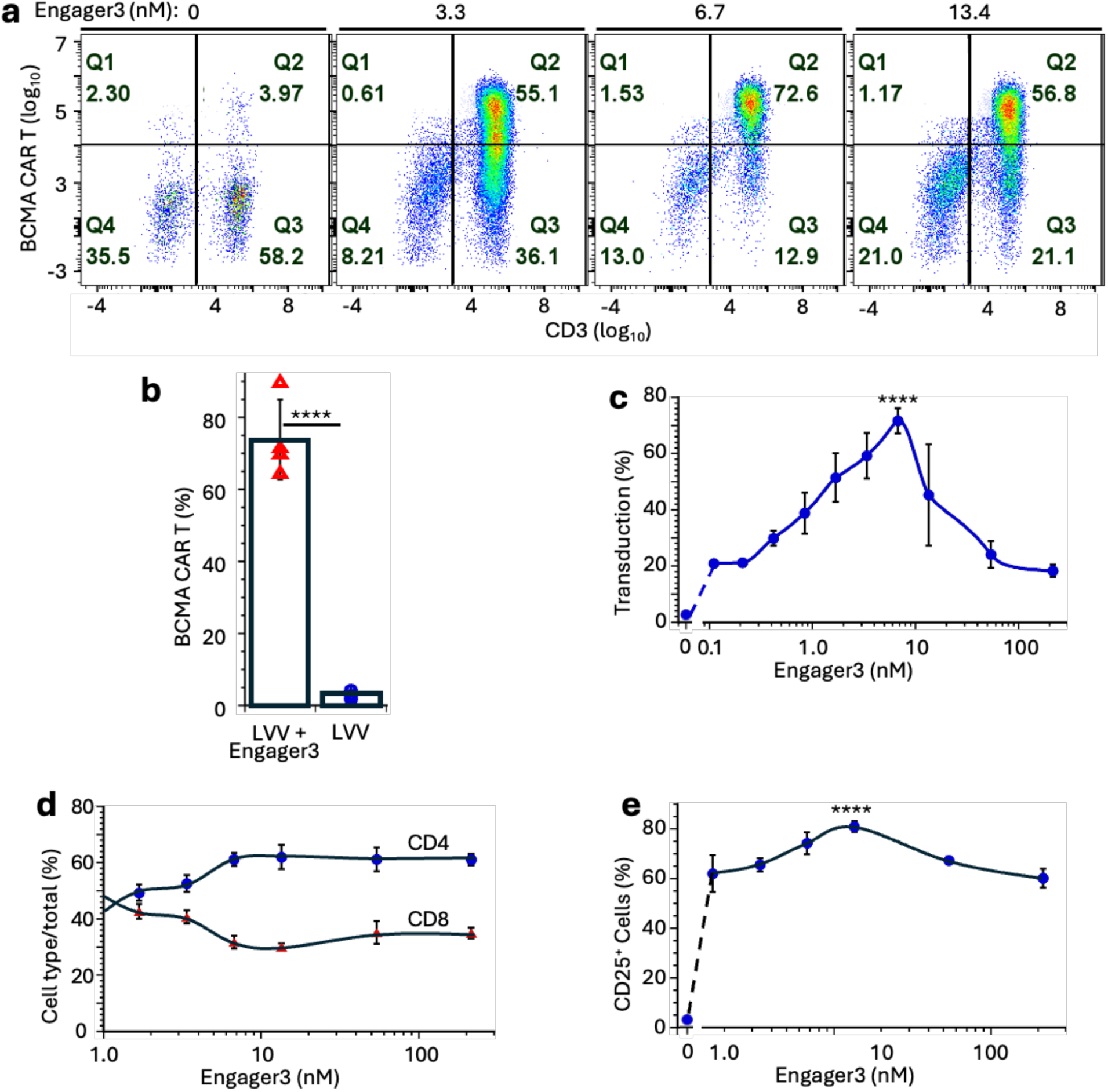
Trimeric engager-mediated transduction generates BCMA CAR T cells in PBMCs. **a,** Representative flow cytometry analysis of BCMA CAR expression in PBMCs. Values indicate the percentage of cells in each quadrant. See **b** for details. **b**, Flow cytometry summary of BCMA CAR expression 7 days following transduction of PBMC (4 donors) by LVV in the presence or absence of Engager3 (7 nM). T cells were gated as CD45-positive, CD3-positive cells. Data in **b** are the mean ± SEM of biological replicates of PBMC from 4 donors. Statistical significance was assessed by one-way ANOVA with Tukey’s Post HOC test against the no-engager control. n = 4; ****P < 0.0002. **c,** Percentage of BCMA CAR-positive T cells after transduction with BCMA CAR-encoding LVV pre-incubated with increasing concentrations of the trimeric Engager3. T cells were gated as CD45-positive, CD3-positive cells. **d,** Distribution of CD4-positive and CD8-positive subsets within the CD45-positive, CD3-positive population. **e,** Expression of CD25 in CD45-positive, CD3-positive T cells seven days after transduction. Data in **c** to **e** are mean ± SEM of technical replicates from a representative donor. Statistical significance was assessed by one-way ANOVA with Dunnett’s multiple-comparisons test against the no-engager control. For **c–e**, data are mean ± SEM of three technical replicates. ****P < 0.0001. For all calculations, the molarity of the trimeric engager is based on its monomeric subunit.

### Engager-mediated LVV generates functional BCMA CAR T cells

The functionality of BCMA-CAR T cells generated from PBMCs in the presence of the trimeric Engager3 was subsequently assessed by measuring their cytotoxic activity against BCMA-expressing tumor cells. Human PBMCs were transduced with BCMA-CAR-encoding LVV, alone or pre-mixed with the trimeric engager at 9 nM. Live cell counts were determined at day 10 post-transduction. The trimeric engager greatly increased the live cell counts as compared with those in cultures incubated without the engager (Fig. 8a). Next, transduced effector PBMCs were co-cultured with luciferase-expressing OPM2-Fluc and U266B-Fluc target cells at several E:T ratios. PBMCs transduced by BCMA-CAR-encoding LVV in the presence of the trimeric Engager3 demonstrated robust cytotoxic activity against these BCMA-expressing tumor cells, resulting in a substantial reduction in tumor cell signal compared with PBMCs transduced without the engager (Fig. 8b, c). These results demonstrate that combining LVV with the trimeric engager enables the effective generation of functional BCMA-CAR T cells in PBMCs, capable of antigen-specific tumor cell killing *in vitro*.

**Fig. 8.**
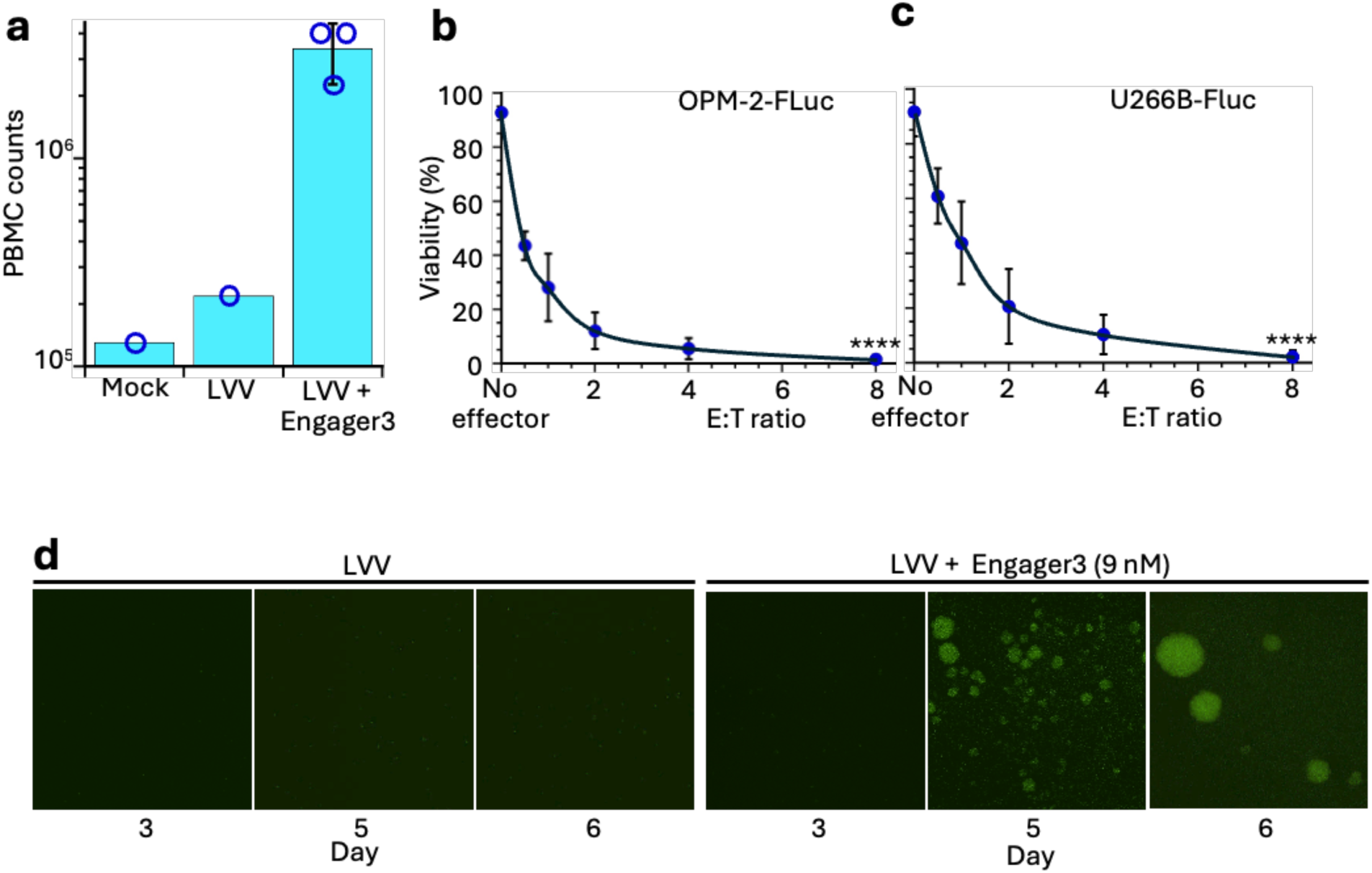
Trimeric engager-generated BCMA CAR T cells are functional. **a,** PBMC counts on day 10 after the transduction with BCMA CAR-encoding LVV alone or pre-incubated with the trimeric Engager3. **b, c,** Transduced PBMCs were co-cultured with OPM2-Fluc (**b**) or U266B-Fluc (**c**) target cells at the indicated effector-to-target ratios. Tumor-cell viability was assessed after 24 h using luminescence relative to time = 0. Viability of OPM2-Fluc cells incubated with effector cells only at E:T = 8 was 76 %. Viability of OPM2-Fluc cells incubated with effector cells transduced with BCMA CAR-encoding LVV without engagers was 37%. Corresponding values for U266B-Fluc cells were 70% and 47%. Data are the mean ± SEM of three technical replicates. Statistical significance was assessed relative to untreated target cells within each condition using one-way ANOVA with Dunnett’s multiple comparisons test. ****P < 0.0001. **d,** Transduction in whole human blood. Fresh blood was incubated with EGFP-encoding LVV alone or LVV preincubated with trimeric engager (final concentration, 9 nM). PBMCs were isolated after 6 h, and EGFP expression was imaged on the indicated days.

To further test the potential of the trimeric Engager3 for in vivo applications, we evaluated its ability to mediate T-cell transduction in human blood. EGFP-encoding LVV was pre-incubated, either alone or with the trimeric engager, and then added to whole human blood. After 6 hours, PBMCs were isolated, and EGFP expression was assessed. Samples transduced by LVV without the engager did not express EGFP. In contrast, the trimeric engager enabled LVV transduction, as evidenced by fluorescence imaging on days 5 and 6 (Fig. 8d).

### The engager-LVV complex enables in vivo CAR T cell generation and subsequent sustained suppression of detectable tumor burden

To evaluate the efficacy of retargeted lentiviral vectors for generating BCMA CAR T cells in vivo, multiple myeloma tumors were established in NSG mice by intravenous administration of OPM-2-Fluc cells on Day -21. On Day -8, human PBMCs were administered intraperitoneally. On day −1, baseline tumor burden was measured by bioluminescence imaging, and mice were assigned to four treatment groups based on tumor radiance. On day 0, a control Saline group received saline IV; an Engager-only group received the trimeric Engager3 IV; an LVV group received LVV IV; and the LVV + Engager group received LVV pre-mixed with the trimeric Engager3 IV. Tumor progression was monitored longitudinally by bioluminescence imaging. Control and Engager-only groups showed rapid tumor progression, confirming aggressive tumor growth in untreated animals. In contrast, the LVV + Engager3 group showed a rapid decline in tumor signals that remained near background levels after four weeks, resulting in sustained survival in all mice (Fig. 9a, Supplementary Figure 5a). The LVV-only group of mice exhibited measurable tumor control, indicating that systemic delivery of lentiviral particles can elicit anti-tumor activity in this mode. However, non-specific transduction by free LVV, particularly in tumor cells, is a risk factor. Circulating CAR T-positive cells were detected in the blood of the LVV group on days 15, 23, and 30 post-transduction, with higher percentages in the LVV + engager group (Supplementary Fig. 5b). All 9 mice in the LVV + Engager group regained body weight, consistent with improved clinical condition. In contrast, three of the 5 mice in the Saline group exhibited severe weight loss and eventually perished. (Supplementary Fig. 5c).

**Fig. 9.**
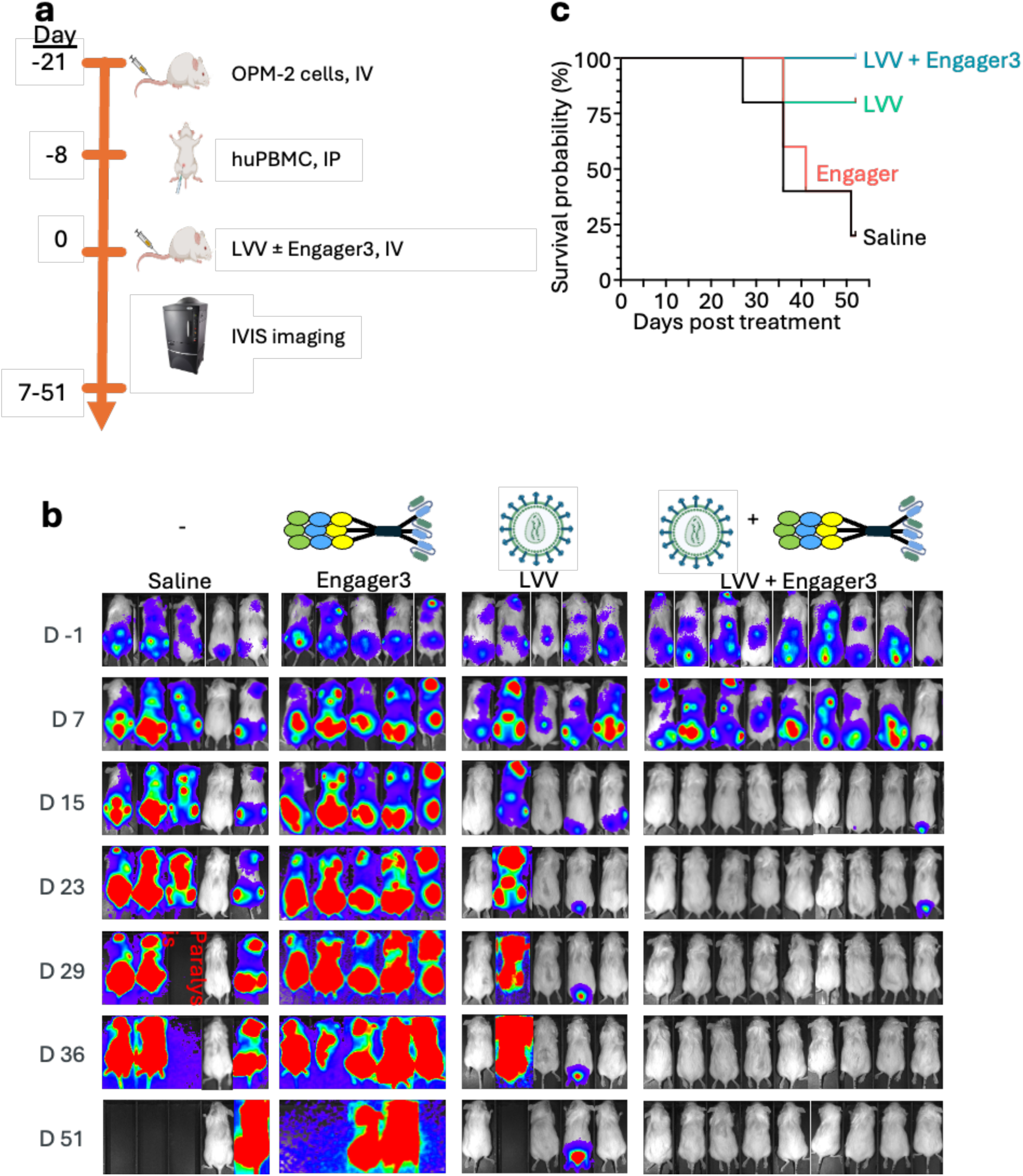
In vivo trimeric engager-mediated CAR T-cell generation controls myeloma. **a,** Study design. Multiple myeloma xenografts were established in NSG mice by IV administration of OPM-2-Fluc cells (2e6 cells/mouse) on Day -21. On Day -8, human PBMC were administered IP. On Day 0, mice received IV saline (5 mice), trimeric Engager3 alone (0.83 µg per mouse, 5 mice), BCMA CAR-encoding LVV alone (1e9 particles/mouse, 5 mice), or a preformed LVV + the Engager3 complex (9 mice). **b,** Longitudinal IVIS imaging of tumor burden in all mice for 51 days. **c**, Kaplan–Meier survival curves of the treated animals. Survival was monitored over the study period and plotted as probability of survival versus days post-treatment. Statistical significance of the LVV + Engager3 group relative to the Saline group using Log-rank (Mantel-Cox) was P < 0.0015.

Together, these results demonstrate that trimeric-engager-coated LVVs enable in vivo CAR T cell generation, resulting in sustained tumor control and prolonged survival in tumor-bearing mice.

## Discussion

In this study, we describe a general platform for redirecting vectors to specific cell types, enabling in vivo CAR T cell therapy and improving oncolytic virotherapy. Specifically, we introduce reversible bispecific engagers based on VSV-G binding domains derived from LDLR fused to ligands of cell-specific receptors. These engagers enable the retargeting of VSV-G– expressing vectors to defined cell types. A major advantage of this approach is that it provides high target-cell specificity while preserving the potency and stability of unmodified VSV-G– based vectors.

Current lentiviral vector (LVV)-based in vivo transduction of specific cell types typically relies on a mutated VSV-G on the vector envelope to abrogate nonspecific entry mediated by the broadly expressed low-density lipoprotein receptor (LDLR) and its family members. However, such mutations are insufficient to completely prevent vector binding to LDLR^32^. Furthermore, they frequently compromise transduction efficiency compared with vectors pseudotyped with unmodified VSV-G^33^. In contrast, the engagers described here effectively suppress nonspecific LDLR-mediated transduction under the conditions tested while maintaining efficient transduction of target cells. Recently, a bispecific adapter consisting of a VSV-G nanobody fused to an ERBB2-binding DARPin was used to direct a lentiviral vector to ERBB2-expressing tumor cells. However, that approach did not eliminate the need to mutate the VSV-G spike to avoid nonspecific transduction^34^.

The trimeric versions of these engagers described here further increase binding avidity, with the three arms of the engager binding simultaneously to the native trimeric VSV-G spike. These multivalent interactions further increased the potency of the engagers as inhibitors of nonspecific, LDLR-mediated entry (Figs. 3 and 6). Following internalization, the rapid vector release from the engager at low endosomal Ca^2+^ recapitulates the natural dissociation of VSV-G from LDLR, resulting in similarly high vector fusion efficiency and gene delivery through the endosomal membrane. Thus, unlike approaches that modify VSV-G to reduce its native tropism, the engager serves as a reversible external targeting component that blocks nonspecific entry before internalization and releases unmodified VSV-G within the endosome.

This feature is particularly important for CAR T cell generation in vivo. This modality is being actively explored as an alternative to approved ex vivo CAR T cell therapies to reduce manufacturing complexity, cost, and treatment-associated risks. The transduction efficiency of non-activated T cells in PBMCs is typically less than 10%^10, 35^. In comparison, at 10 nM, transduction of T cells in PBMCs without prior TCR activation using Engager3 achieved ∼60-70% efficiency (Fig. 7). Efficiency is a critical parameter given the cascading attrition in in vivo CAR T cell therapy^36^. Furthermore, the engager enables efficient transduction without prior T-cell activation, a feature that may help preserve T-cell stemness^37^. Indeed, effective clearance of OPM-2 multiple myeloma in the NSG mouse model was achieved using 0.83 μg of a 46.3 kDa Engager3 per mouse, corresponding to calculated initial engager concentration of 10 nM in ∼2 mL of mouse blood (Fig. 9). This concentration of the trimeric Engager3 effectively inhibited nonspecific transduction of tumor B cells and TCR-deficient Jurkat T cells (Fig. 6). Free LVV also generated detectable circulating BCMA CAR T cells and produced partial tumor control (Fig. 9b and Supplementary Fig. 5a). Because uncoated LVV retains LDLR-dependent tropism, it may also transduce proliferating non-target cells, as illustrated by the transduction of CD3-negative NALM6 cells (Fig. 6e). Thus, this engager provides an important advantage by directing LVV toward T cells while simultaneously limiting LDLR-mediated transduction of non-target cells.

The use of engager-coated unmodified LVV offers additional practical advantages. Vectors pseudotyped with unmodified VSV-G are relatively stable, can be concentrated to high titers, and are supported by well-established large-scale manufacturing and handling protocols. By contrast, vectors based on modified VSV-G often require extensive re-engineering, with unpredictable consequences for yield and storage stability. Vector-Engager3 complexes maintained functional integrity across multiple freeze-thaw cycles and extended room-temperature exposure, a critical logistical attribute for scalable, commercially viable LVV manufacturing. Although our standard protocols include a 1-hour pre-incubation of the trimeric engager with the vectors, comprehensive kinetic analyses revealed that the binding equilibrium is established within one minute (data not shown). Together, these properties suggest that target specificity can be added to an established VSV-G-pseudotyped LVV platform without sacrificing its favorable transduction, manufacturing, and handling properties.

This retargeting approach is also directly applicable to the use of VSV as an oncolytic virus. Current clinical trials employ attenuated VSV variants to avoid neurotropism. Our finding that CEACAM8-based engagers can specifically infect CEACAM6-overexpressing tumor cells with VSV, but not CEACAM6-negative cells, suggests a strategy to improve both the safety and efficacy of VSV-based oncolytic therapies. Confining VSV tropism to tumor tissue may lower the viral dose required for therapeutic activity by preventing virus sequestration in healthy, VSV-resistant tissues. Moreover, even if certain healthy cells remain permissive to VSV uptake, administration of excess high-affinity trimeric CEACAM8-(CR2)₃ engager may effectively block nonspecific infection by both input virus and progeny virions (Fig. 3c). In contrast, infection of CEACAM6-positive cells was not inhibited even at micromolar concentrations of excess engager (Fig. 3d). The effective cytolysis of target tumor cells at high CEACAM8-(CR2)₃ engager concentrations indicates that the engager-vector interaction is fully reversed within the endosome. Thus, in contrast to incomplete restriction of viral entry by modifying the viral glycoprotein, the reversible engager may provide systemic protection against nonspecific infection while preserving VSV’s full infectivity after uptake by the targeted tumor cell.

Importantly, CEACAM8-(CR2)₃ consists exclusively of human receptor-derived domains. Therefore, even high systemic concentrations are unlikely to elicit adverse immune responses. CEACAM8-based engagers may therefore have therapeutic potential in breast, pancreatic, colorectal, and ovarian cancers, which frequently overexpress CEACAM6. CAR T cell therapy is largely limited to hematopoietic malignancies. Recently, combined treatment with CAR T cells and oncolytic VSV was found to be highly effective in a melanoma model in mice, possibly expanding the success of CAR T therapy into solid tumors^19^. The ability of the same engager platform to support both targeted in vivo CAR T cell generation and tumor-directed oncolytic VSV may therefore provide complementary approaches to cancer therapy.

Beyond these two applications, our finding that CEACAM8-based engagers enable selective transduction of CEACAM6-expressing epithelial cells suggests that these engagers could facilitate luminal delivery of lentiviral vectors to epithelial cells in the respiratory and gastrointestinal tracts. By swapping the targeting domains, this technology may enable reprogramming of diverse cell types, including natural killer cells, monocytes, dendritic cells, and hematopoietic stem/progenitor cells, thereby establishing a versatile therapeutic system for in vivo gene editing and immunotherapy. Similarly, VSV-G-based virus-like particles^38^ and non-integrating lentiviral vectors coated with targeting engagers may support cell-type-specific CRISPR-based gene-editing strategies. Furthermore, the trimeric architecture readily accommodates the multiplexing of heterologous ligands on a single viral particle, a capability particularly valuable for oncology applications.

Collectively, our findings position the engager-based platform as a scalable and efficient strategy for targeted in vivo gene delivery, gene editing, immunotherapy, and oncolytic virotherapy without sacrificing the favorable properties of VSV and unmodified VSV-G-pseudotyped vectors. The ability to separate cell targeting from the viral vector itself, while preserving the native activity of VSV-G, represents a central advantage of this approach and may facilitate the development of a common vector platform that can be redirected to different cell types by exchanging the targeting component of the engager.

## Methods

### Cells and vectors

Human FS11 foreskin fibroblasts^29^ were kindly provided by M. Revel. All other cell lines were from ATCC. The TCR knockout and LDLR knockout lines of Jurkat were generated in-house. Human peripheral blood mononuclear cells (PBMCs) were obtained from healthy donors and isolated by density gradient centrifugation using Ficoll-Paque PLUS (171440, Cytiva). Cells were washed and resuspended in ATCC-modified RPMI-1640 medium (ATCC RPMI, 30-2001, ATCC) supplemented with 10% fetal bovine serum (FBS) (A52567-01, Gibco), 1× GlutaMAX (35050061, Gibco), and recombinant human IL-2 (25 ng/mL) (200-02, PeproTech). BCMA-positive OPM-2 and U266B cell lines were engineered to express the firefly luciferase gene. The VSV Indiana strain, obtained from ATCC, was propagated in Vero cells. The EGFP- and mCherry-encoding lentiviral vector (pLV[Exp]-EGFP/Puro-EF1A>mCherry) is from VectorBuilder. The VSV-G-pseudotyped lentiviral vector encodes either the SFFV promoter driving EGFP expression or the LCK promoter driving the BCMA-CAR transgene in reverse orientation (Imanis Life Sciences). The BCMA-CAR expression cassette is driven by an internal T-cell selective promoter-enhancer element consisting of an LCK promoter linked to an mCMV enhancer.

### Production and purification of recombinant proteins

We designed mammalian expression vectors encoding the following engagers:

(1). sLDLR. A control protein corresponding to human pro-LDLR_(25-145)_ (GenBank accession No. NP_000518.1), followed by a His tag (GTKHHHHHH). It was constructed in a Mammalian expression vector and expressed in CHO-S cells. The protein was purified using a Ni-NTA column followed by size-exclusion chromatography (GenScript).

(2). sLDLR-CEACAM8. This engager consists of human pro-LDLR_(25-145)_, a linker (GSGGGGSGG), human pro-CEACAM8_(35-140)_ (GenBank accession No. NP_001807.2), and a His_6_ tag. It was constructed in the pcDNA3.1 vector and expressed in HEK293T cells. The protein was purified using a Ni-NTA column. This engager was used in experiments described in Figs. 2a, b, and 4.

(3). CEACAM8-sLDLR. This engager consists of CEACAM8_(35-140)_, a 9-amino acid residues linker, sLDLR_(25-145)_, and a His tag. It was constructed in a Mammalian expression vector and expressed in CHO-S cells. The protein was purified using a Ni-NTA column followed by size-exclusion chromatography (GenScript). This engager was used in experiments described in Figs. 2c, 3, and Supplementary Fig. 3.

(4). sCEACAM6. Human proCEACAM6 _(35-140)_ fused to a His_6_ tag was produced in CHO-S cells and purified using a Ni-NTA column and size-exclusion chromatography (GenScript).

(5). CEACAM8-(CR2)_3_. This trimeric engager had the general formula CEACAM8_(35-140)_-L1-CR2-L2-CR2-L2-CR2-His_6_, where L1 is a 10-amino-acid linker, and L2 is 34-amino-acid-long. It was produced in CHO-S cells and purified using a Ni-NTA column followed by size-exclusion chromatography (GenScript).

(6). Engager3. This engager consisted of sLDLR_(25-145)_, a (G_4_S)_3_ linker, the trimerizing *VL* peptide AAQPAMKQLEDKVEELLSKVYHLENEVARLKKLVGE, corresponding to the leucine zipper domain of yeast GCN4 with Val and Leu in the a and d positions of the heptad repeats, respectively ^39, 40^, a G_4_S linker, UCHT1 (a CD3-targeting scFv ^41^), a GSGGSG linker, and a FLAG tag. It was cloned into the pcDNA3.4 vector, produced in CHO-S cells, and purified using Protein-L purification columns followed by size-exclusion chromatography (GenScript). All reported concentration values are based on the monomeric form of this engager (46.3 kDa).

1. Engager1. This monomeric engager is identical to Engager3 except that the *VL* sequence was replaced with the non-trimerizing peptide AAQPAMKQAEDKAEEALSKAYHAENEAARAKKLVGE, having the same length as *VL* and Ala substitutions in the a and d positions of the heptad repeats.

For BLI experiments, both Engager3 and Engager1 were produced using Viral Production Cells 2.0 (VPC 2.0; Gibco) and purified using Protein-L purification columns and size-exclusion chromatography (Vyriad).

### Assay of VSV infectivity in the presence or absence of engagers

AsPC-1 cells (100,000/well), in 0.1 mL of RPMI-1640 medium supplemented with penicillin (100 IU/mL), streptomycin 0.1 mg/mL, sodium pyruvate (1 mM), glutamine (2 mM), and 10% Fetal bovine serum (RPMI-10) were seeded in 96-well plates and incubated for 2 h at 37°C in 5% CO_2_. VSV was diluted separately in RPMI-10 to 6.25 × 10^4^ PFU/mL, and 0.1 mL aliquots in 96-well plates were pre-incubated for 15 min at room temperature with twofold serial dilutions of sLDLR-CEACAM8 or sLDLR_25-145_, starting at 1 μM. The VSV-engager mixtures were then added to confluent AsPC-1 cell cultures. The cultures were further incubated at 37 °C in 5% CO_2_ for 24 hours. The media were then discarded, and the cells were fixed for 15 minutes with cold methanol. They were then stained with 2.3% Crystal violet (HT90132, Sigma-Aldrich) diluted 1:5 in 70% ethanol. The plates were then washed twice with ddH_2_O, and cell viability was measured using a TECAN Infinite M Plex plate reader at 590 nm. Control, untreated AsPC-1 cell cultures and cultures treated with VSV alone were used as standards of 100% and 0% protection, respectively. Assay of VSV infectivity in HeLa cells was similarly performed except that 12,000 cells/well were seeded in 96-well plates in DMEM-10 (DMEM supplemented with 100 IU/mL penicillin, 0.1 mg/mL streptomycin, 2 mM glutamine, and 10% FBS). Assay in FS11 fibroblasts was performed as in HeLa cells, except that the cell density was 20,000 cells/well.

### Transduction of adherent cells

AsPC-1 cells (100,000 cells/well) in RPMI-10 (0.1 mL), Caco-2 cells (50,000 cells/well) in DMEM 20 (0.1 mL), and FS11 fibroblasts (20,000 cells/well) in DMEM-10 (0.1 mL) were seeded in black 96-well plates (Greiner) and incubated for 2 hours at 37°C and 5% CO_2_. Stock lentiviral vector encoding EGFP (6x10^9^ TU/ml) was pre-incubated with the indicated engager concentrations in Opti-MEM for 15 min. at room temperature. The media was removed from the cell cultures, and 30 μL of lentiviral vector-engager mixture, corresponding to 2 μL of LVV stock, was added to the cells for 1 hour. The corresponding media (170 µL) was then added, and the incubation continued for 24 hours. All media were then replaced with fresh media (0.2 mL), and the cultures were further incubated for an additional 48 hours. The media were then replaced with PBS, and fluorescence was measured using the Infinite 200 PRO plate reader with excitation at 485 nm and emission at 515 nm. GFP-expressing cells were also photographed by a Nikon Eclipse Ti fluorescent microscope.

For experiments involving CEACAM8-(CR2)_3,_ we used an LVV stock encoding both EGFP and mCherry. The incubation time of engagers with LVV was extended to 1 hour, and mCherry fluorescence was measured with excitation at 580 nm and emission at 610 nm using a TECAN Infinite M Plex plate reader.

### Competition assay

AsPC-1 pancreatic adenocarcinoma epithelial cells (100,000 cells/well in 100 μL RPMI-10) were seeded in 96-well plates and left to adhere for 2 h at 37°C. VSV (6.25 × 10^4^ PFU/mL) was mixed with the specified concentrations of sLDLR, CEACAM8-sLDLR, and the competitor sCEACAM6. The mixtures were allowed to stand at room temperature for 15 minutes. The AsPC-1 culture media were replaced with 200 μL of the engager and competitor mixture, and incubation continued at 37°C for 24 hours. The media were then discarded, and the cells were fixed for 15 minutes with cold methanol. They were then stained with 2.3% Crystal violet (Sigma HT90132) diluted 1:5 in 70% ethanol. The plates were then washed twice with ddH_2_O, and the cell viability was measured using a TECAN Infinite M Plex plate reader at 590 nm.

### Immunostaining of CEACAM6

Cells (AsPC-1, FS11, Caco-2, HeLa) were grown to confluency in ibidi 8-well µ-Slides, then washed with PBS and fixed with 4% paraformaldehyde. The plates were blocked with 10% horse serum for 1 hour and then incubated with mouse anti-human CEACAM6 (1:50, sc-69899, Santa Cruz) in 3% BSA in PBS for 2 hours. The plates were then washed and incubated for 1.5 hours with Cy3-labeled donkey anti-mouse Ig antibody (1:100, AP192C, Sigma-Aldrich). Cell nuclei were stained with DAPI (1:2000) for 1 minute, and the plates were washed with ddH_2_O. The cultures were then photographed by a Nikon Eclipse Ti fluorescent microscope

### Biolayer Interferometry (BLI)

Recombinant Human CD3E & CD3G heterodimer receptor (Avi-tag Biotinylated/His tag; CT123-H49H-B, Sino Biological) was immobilized to Flex SA sensors (350001, GatorBio) at 10 μg/mL in a Binding buffer (20 mM HEPES, pH 7.4, 150 mM NaCl, 2 mM CaCl_2_). Kinetics assays were carried out at 30°C using a standard acquisition rate of 5.0 Hz, averaging 20 readings, with a sample plate shake speed of 1000 rpm. After baseline measurement in the Binding buffer alone, the binding kinetics were monitored in the Association step by dipping the biosensors in wells containing the designed proteins in 1:3 serial dilutions (6.83 µM to 8.8 nM for the trimeric Engager3, and 20.5 µM to 26.6 nM for the monomeric Engager1). The sensors were then dipped into a dissociation buffer (20 mM HEPES, pH 7.4, 150 mM NaCl) in 96-well plates. Binding data were collected in a Gator Prime (Gator Bio) and processed using the instrument’s integrated software (GatorOne v2.18). For the monomeric engager, the binding curves were analyzed using a formal 1:1 Langmuir Binding model, reporting absolute *Kd* values. Due to the trivalent nature of the trimeric engager and the assay format used (with avidity), the *Kd* values determined for the trimeric engager are considered apparent *Kd*.

### Transduction of suspension cells

Suspensions of Jurkat cells, Jurkat-TCR-KO cells, and Nalm6 cells were seeded at 5e4 cells/well in 96-well plates in 0.1 ml of ATCC RPMI containing 10% FBS (ATCC RPMI-10). For measuring IC_50_, EGFP-encoding LVV (2.5e8 particles based on p24), alone or mixed with either the trimeric Engager3 (7.7-153.8 nM) or the control monomeric Engager1 (0.081 to 3.85 µM), was incubated for 1 hour at room temperature. Then, 15 µl reaction mixtures were added onto the Jurkat or Jurkat-TCR-KO cell cultures, giving final concentrations of 1 to 20 nM of the trimeric engager and 10 to 500 nM of the monomeric engager. The plates were incubated for 3 days at 37°C, and the percentage of EGFP-expressing cells was determined by flow cytometry. Images were captured using a Nikon 10x fluorescent microscope. For molar calculations, the molecular mass of the monomeric subunit of Engager3 was used.

### Transduction of PBMCs with BCMA-CAR encoding LVV

Human peripheral blood mononuclear cells (PBMCs, leukopak) were thawed in 6 mL of 50% FBS in ATCC RPMI and centrifuged (350 × g, 5 min). The cell pellet was resuspended in 5 mL of complete medium (CM; ATCC RPMI-10 and 25 ng/mL of human IL-2) and counted using a CellDrop (DeNovix Inc.) with 0.4% trypan blue viability staining (T8154, Sigma-Aldrich). Cells were adjusted to 3x10⁶ cells/mL in CM and cultured overnight in T25 flasks at 37°C with 5% CO₂ to allow recovery from thawing. The next day (Day 0), PBMCs were washed once, resuspended at 4 × 10⁶ cells/mL in fresh CM, and placed in U-bottomed wells at 2 × 10⁵ cells/well (50 μL/well). BCMA-CAR-encoding LVV in ATCC RPMI-10 (1000 particles/cell, based on p24 ELISA) was pre-incubated for 1 hour at room temperature, either alone or with various Engager3 concentrations, and then added to the PBMCs cultures at 50 µL/well, bringing the final volume to 100 μL. Final engager concentrations ranged from 0.11 to 215 nM. Mock-infected wells received media alone. The plates were sealed, centrifuged at 800 × g for 30 min at 32°C (spinoculation), and then transferred back to the 37°C incubator. The next day, 100 µl fresh CM was added to each well. Thereafter, half of the media was replaced with fresh CM every other day starting from day 3, and then BCMA-CAR expression was measured on day 7.

For the cell-killing assay, PBMCs were suspended at 5×10⁵ cells per well (250 µL/well) in 12-well plates, and the LVV ± Engager3 was added at 500 µL/well, bringing the total volume to 750 µL/well and 9 nM of Engager3 (molarity was based on its monomeric subunit). The plates were sealed, centrifuged at 800 × g for 30 min at 32°C (spinoculation), and then transferred back to the 37°C incubator. The next day, 250 µL of CM was added to each well, bringing the final volume to 1 mL. Thereafter, half of the medium was replaced with fresh CM every other day starting from day 3, and the cells were counted and harvested for the downstream killing assay on day 10 post-transduction.

### Transduction of whole blood

Fresh blood (1-1.5 mL/well) was placed in a 6-well plate. EGFP-encoding LVV (4e9 particles/well) was mixed with or without the Engager3 for 1 hour at room temperature, then added to the blood (final engager concentration: 9 nM). The blood + LVV ± engager was incubated at 37°C with shaking for 6 hours. PBMCs were then isolated using Ficoll-Paque PLUS gradient centrifugation and cultured at 37°C for the indicated times. EGFP expression was photographed on the indicated days.

### Flow cytometry of hematopoietic cells

Flow cytometry was used to assess CAR expression, T-cell activation, and T-cell subset composition. Cells were harvested and stained with fluorophore-conjugated antibodies against human CD45-FITC (Clone HI30, 304006, BioLegend), CD3-BV605 (Clone OKT3, 317322, BioLegend), CD4-PerCp-Cy5.5 (Clone SK3, 344608, BioLegend), CD8-Pe-Cy7(Clone: SK1, 344750, BioLegend), CD25-BV421(Clone M-A251, 356114, BioLegend), and CD69-BV650 (Clone FN50, 563835, BD Biosciences). BCMA CAR expression was detected using a two-step staining approach, in which CAR binding was detected using Human BCMA/TNFRSF17 protein, Llama IgG2b Fc tag (BCA-H5259, ACROBiosystems), followed by secondary detection using Goat Anti-Llama IgG-AF647 (sAb070, Imanis Life Sciences). Samples were analyzed by flow cytometry (CytoFLEX LX, Beckman Coulter Life Sciences), and the data were analyzed using FlowJo software. CAR expression and activation markers were evaluated within the CD45⁺, CD3⁺ T cell population.

### *In vitro* CAR T cell-mediated killing assays

CAR-transduced PBMCs were cultured for 10 days post-transduction before initiating killing assays. The BCMA-positive multiple myeloma target cell lines OPM2-Fluc and U266B-Fluc (25,000 cells in RPMI-10, 50 μL/well) were placed in 96-well clear-bottomed black plates. Thereafter, the transduced PBMCs effector cells were added to the target cell cultures at several E:T ratios. The cultures were incubated for 24 hours. Bio-Glo^TM^ Luciferase reagent (100 μL/well, G7940, Promega) was added, and the mixture was incubated for 10 minutes. The luminescence was measured using an Infinite 200 Pro-Lumi plate reader (Tecan Life Sciences). Lower luminescence was interpreted as indicating fewer viable tumor cells.

### *In vivo* multiple myeloma model

All animal studies were conducted in accordance with the Mayo Clinic institutional guidelines for the care and use of animals. At day -21, female NOD.Cg-*Prkdc^scid^ Il2rg^tm1Wjl^*/SzJ mice aged 4-5 weeks old (005557, The Jackson Laboratory) were implanted intravenously (IV) with the BCMA^+^ bone-marrow-tropic multiple myeloma cell line OPM2-Fluc (IgG lambda), stably expressing Firefly Luciferase (2 × 10^6^ cells in 200 µL HBSS; CSP305, Imanis Life Sciences). Healthy human PBMC from a single donor were isolated from Leukopack using a Ficoll-Paque PLUS density gradient. Purified PBMCs were frozen in Cryostor CS10 (C2874 Sigma-Aldrich) at 2 × 10^7^ cells per vial and stored in liquid nitrogen until use. On day -9, PBMCs were thawed, washed twice with DPBS, and suspended in ATCC RPMI-10 at 4 x 10^6^ cells/mL. On day -8, PBMCs were washed twice with DPBS, resuspended in HBSS, and 0.2 mL containing 1 × 10^7^ cells was injected into each mouse IP. On day −1, baseline tumor burden was assessed by bioluminescence imaging (BLI), and mice were randomized by body weight and baseline tumor burden into treatment groups via the RandoMice v1.17 program. Mice with insufficient tumor engraftment were excluded. The remaining mice were allocated to an efficacy cohort or a correlative studies cohort based on tumor radiance. On day 0, mice received an IV injection of either saline, the trimeric Engager3 (0.83 μg), BCMA CAR-encoding LVV (1 × 10⁹ particles/mouse, based on DuoSet p24 ELISA), or LVV (1 × 10⁹ particles) pre-mixed for 1 hour with Engager3 (0.83 μg, 89 nM in 0.2 mL). All injected mixtures were brought to a final volume of 0.2 mL. Therapeutic efficacy was assessed by monitoring tumor growth and clearance kinetics using luciferase-based in vivo imaging. The engager dose was 89 nM in the injectate.

### Bioluminescence imaging

Tumor burden was longitudinally assessed by bioluminescence imaging (BLI) using the Xenogen In Vivo Imaging System (IVIS). D-luciferin (15 mg/mL in sterile PBS, 200 μL) was administered IP to the mice 10 minutes before imaging. The animals were then anesthetized with 1–3% vaporized isoflurane, and images were acquired with auto-exposure settings and an f/1 aperture to maximize signal sensitivity.

### Measuring CAR T cells in peripheral blood

Whole blood (50 µL) was collected from each animal in a MiniCollect Tube (0.5 mL K2E K2EDTA, 450532, Greiner Bio-One) on Days 15, 23, and 30 post-transduction. Red blood cells were lysed by adding ACK lysis buffer (150 µl, A1049201, Gibco) to the blood and incubating the samples for 4 minutes. The lysis reaction was terminated by adding 100 μL PBS containing 4% FBS. Samples were then centrifuged at 800 × g for 5 minutes, washed once with 200 µL PBS, and subsequently processed for flow cytometric analysis to measure the percentage of CAR-expressing T cells within the total CD45⁺CD3⁺ T cell population and the T cell subset composition. Cells were harvested and stained with fluorophore-conjugated antibodies against mouse CD45-AF700 (Clone 30F-11, 103128, BioLegend), human CD45-FITC (304006, Clone HI30, BioLegend), CD3-BV605 (317322, Clone OKT3, BioLegend), CD4-PerCp-Cy5.5 (344608, Clone SK3, BioLegend), CD8-Pe-Cy7(344750, Clone SK1, BioLegend), CD25-BV421(356114, Clone M-A251, BioLegend), and CD69-BV650(563835, Clone FN50, BD Biosciences). BCMA CAR expression was detected using a two-step staining approach, in which CAR binding was detected using anti-Llama IgG2b BCMA/TNFRSF17 BCMA antigen (BCA-H5259, ACROBiosystems) followed by secondary detection using Goat Anti-Llama IgG-AF647 (sAb070, Imanis Life Sciences). Samples were analyzed using a flow cytometer (Cytoflex LX, Beckman Coulter), and cell composition was calculated using FlowJo software.

### Statistical analyses

Statistical analyses for the data in Figs 2, 4, 7b, and Supplementary Fig. 3 were performed using ANOVA with Tukey HSD post Hoc test. One-way ANOVA with Dunnett’s multiple comparisons test was used for Fig. 7c. Log-rank (Mantel-Cox) test was used for Fig. 9c. Technical triplicate dose-response curves in Fig. 3c were fitted by nonlinear regression using a four-parameter logistic model. Differences in logIC₅₀ between monomeric and trimeric engagers were evaluated by an extra-sum-of-squares F test.

## Data availability

Excel files associated with the Graphs have been deposited in the Figshare website (https://doi.org/10.6084/m9.figshare.33533224).

## Acknowledgments

This study was supported in part by an internal Grant from the BINA program of the Weizmann Institute of Science. Research related to CD3 engagers was carried out at Imanis Life Sciences, LLC, Vyriad Inc., and Mayo Clinic animal care facility under a research contract from Vyriad, Inc. This contract was funded exclusively by Vyriad. We thank Eric Smith (Dana-Farber Cancer Institute) for providing the BCMA scFv (VH only) sequence.

## Author contributions

G.N.N. designed CD3 engagers, supervised studies, performed data analysis, and assisted in manuscript writing. D.G designed and performed experiments and analyzed data. M.E. performed molecular structure analysis. G.S. supervised studies and evaluated data. M.S. performed CD3 engager complexation experiments and transductions on suspension cells. N.P. designed in vivo experiments. P.J. performed whole-blood PBMC transductions and mouse-related procedures and analyses. T.K. and H.A. performed CD3 engager complexation experiments and transductions on suspension cells. C-H.C. and A.K. performed whole blood PBMC transductions and mouse-related procedures and analysis. H.S., P.L., L.B., and M.A.M.A performed Bio-Layer Interferometry (BLI) assays, CD3 engager purifications, and FLAG-tag ELISA assays. K.W.P provided input on interpreting the results and participated in the data discussion. S.J.R. designed trimeric CD3 engagers and supervised studies. M.R. Conceived the idea, designed studies, supervised the research, and wrote the manuscript.

## Additional information

Supplementary information accompanies this paper.

## Competing interests

M.R., D.G., and G.S. are co-inventors on patent applications. G.N.N., M.S., N.P., P.J., T.K., H.A., A.K., H.S., P.L., L.B., M.A.M.A., K.W.P., and S.J.R. are Company employees.

