## Supplemental data for "Bispecific VSV-G Engagers: A Platform for Cell-Type-Specific Delivery of Viral Vectors"

Supplementary Fig. 1

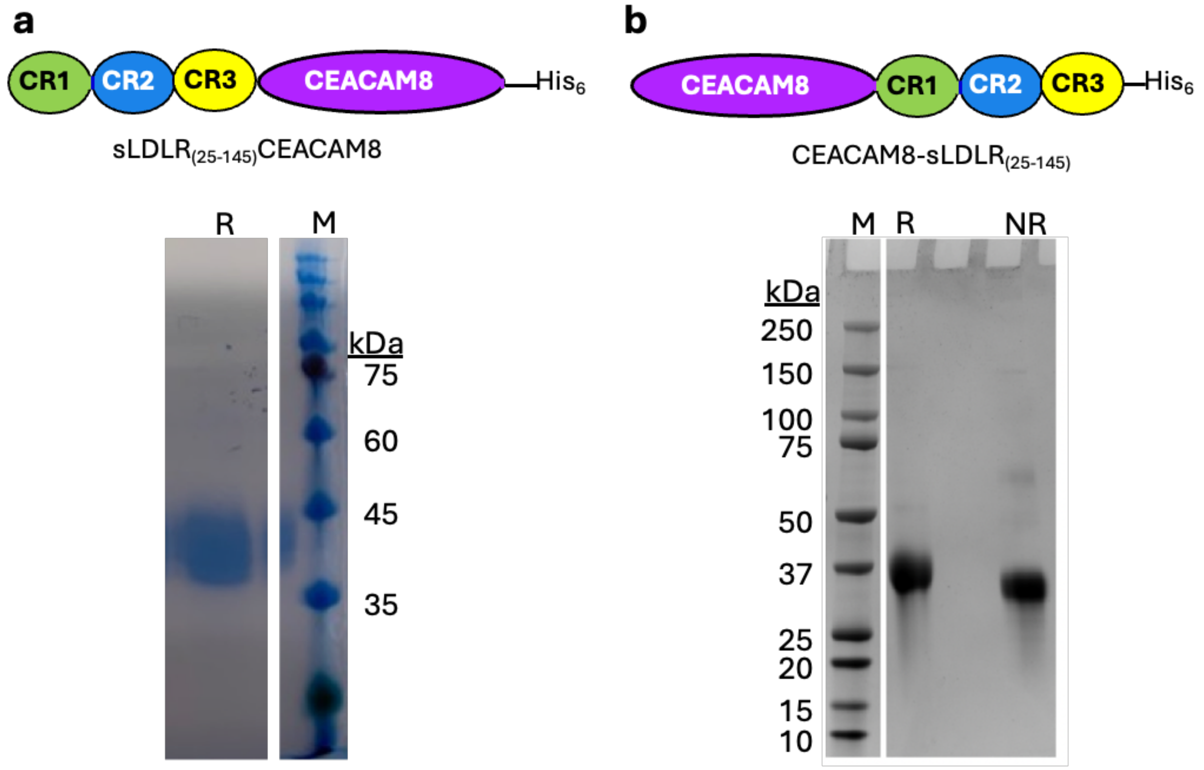


**Supplementary Fig. 1 | Design and production of CEACAM8-based bispecific engagers. a,** sLDLR-CEACAM8 was expressed in HEK293T cells. **b,** CEACAM8-sLDLR was expressed in CHO-S cells. Both proteins were purified by Ni-NTA chromatography and analyzed by SDS-PAGE. M, molecular-mass markers (kDa); R, reducing conditions; NR, non-reducing conditions. Broad bands are attributed to N-linked glycosylation of the CR domains.

Supplementary Fig. 2

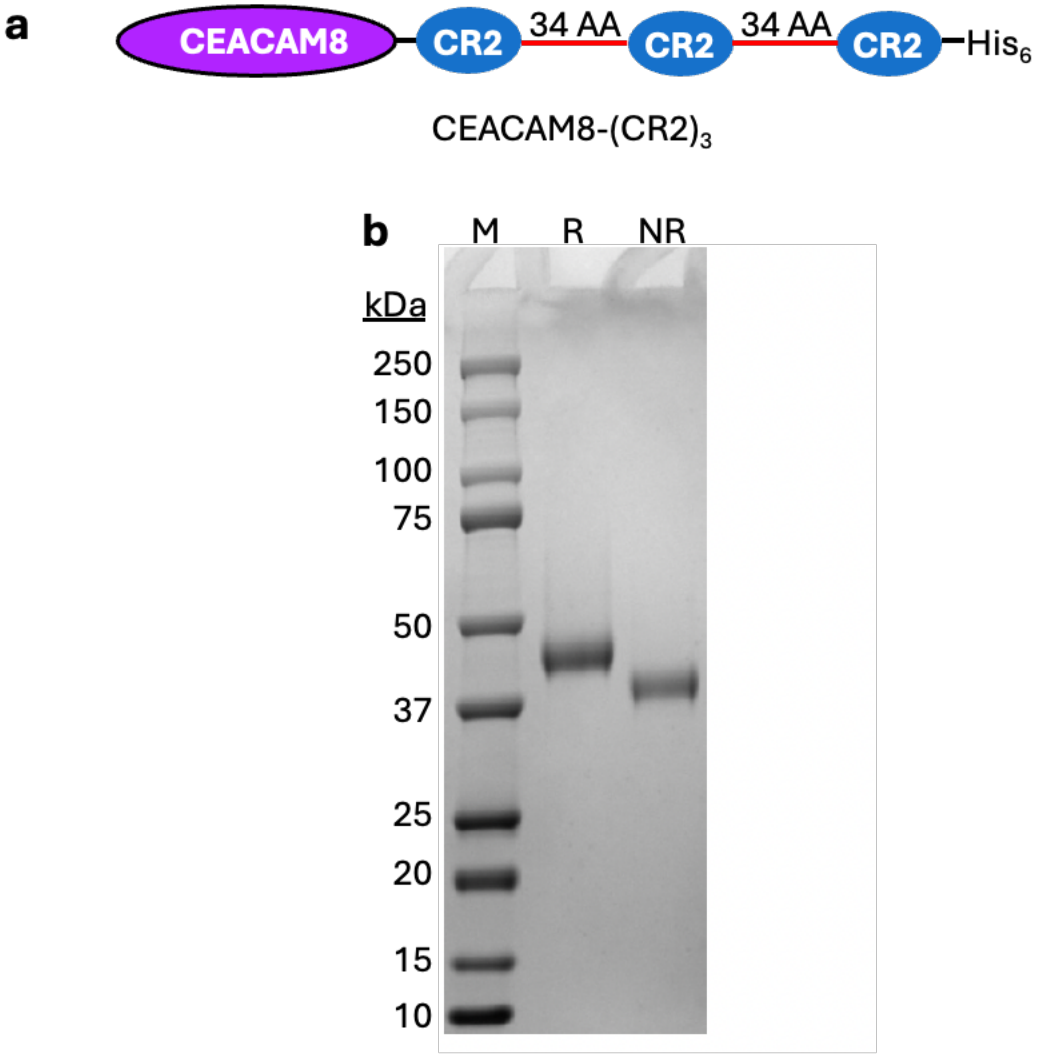


**Supplementary Fig. 2 | Design and purification of CEACAM8-(CR2)$_3$. a,** Schematic representation of the reversible multivalent engager CEACAM8-(CR2)$_3$. The protein was expressed in CHO-S cells and purified by Ni-NTA chromatography. **b,** SDS-PAGE of purified CEACAM8-(CR2)$_3$. M, molecular-mass markers (kDa); R, reducing conditions; NR, non-reducing conditions.

### Supplementary Fig. 3

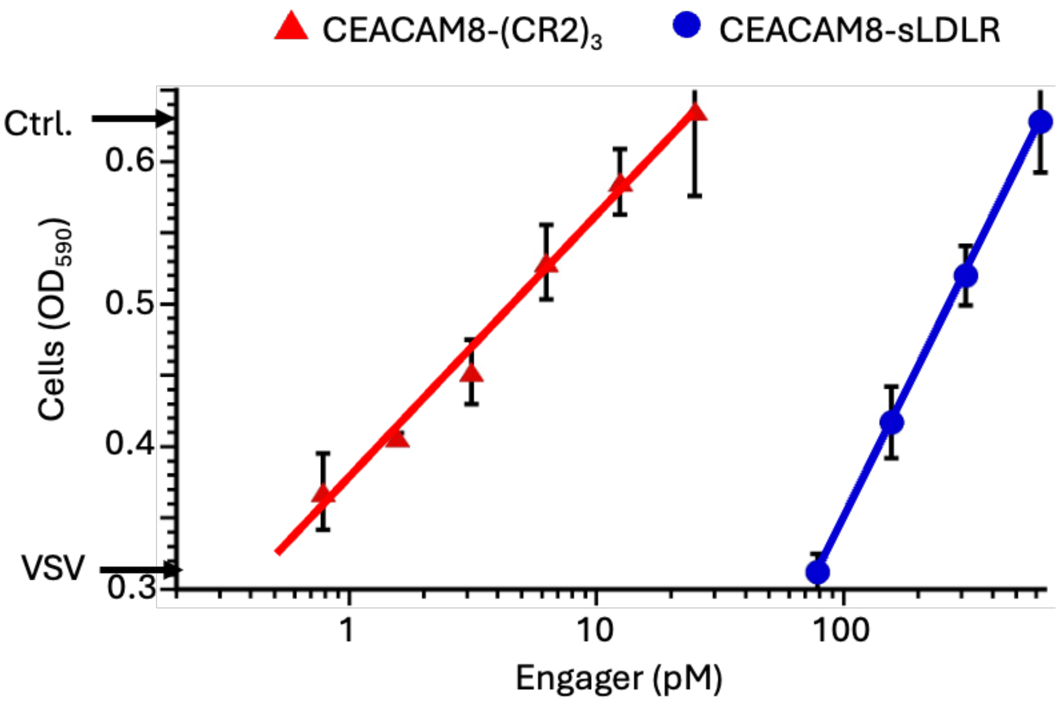


**Supplementary Fig. 3 | CEACAM8–(CR2)3 blocks VSV cytolysis of HeLa cells.** Comparison of CEACAM8-sLDLR and CEACAM8-$(CR2)_3$ as inhibitors of VSV cytolysis in CEACAM6-negative HeLa cells. Experimental conditions were as described in Fig. 3. Arrows on the Y-axis indicate untreated cultures and cultures exposed to VSV alone (n = 4).

Supplementary Fig. 4

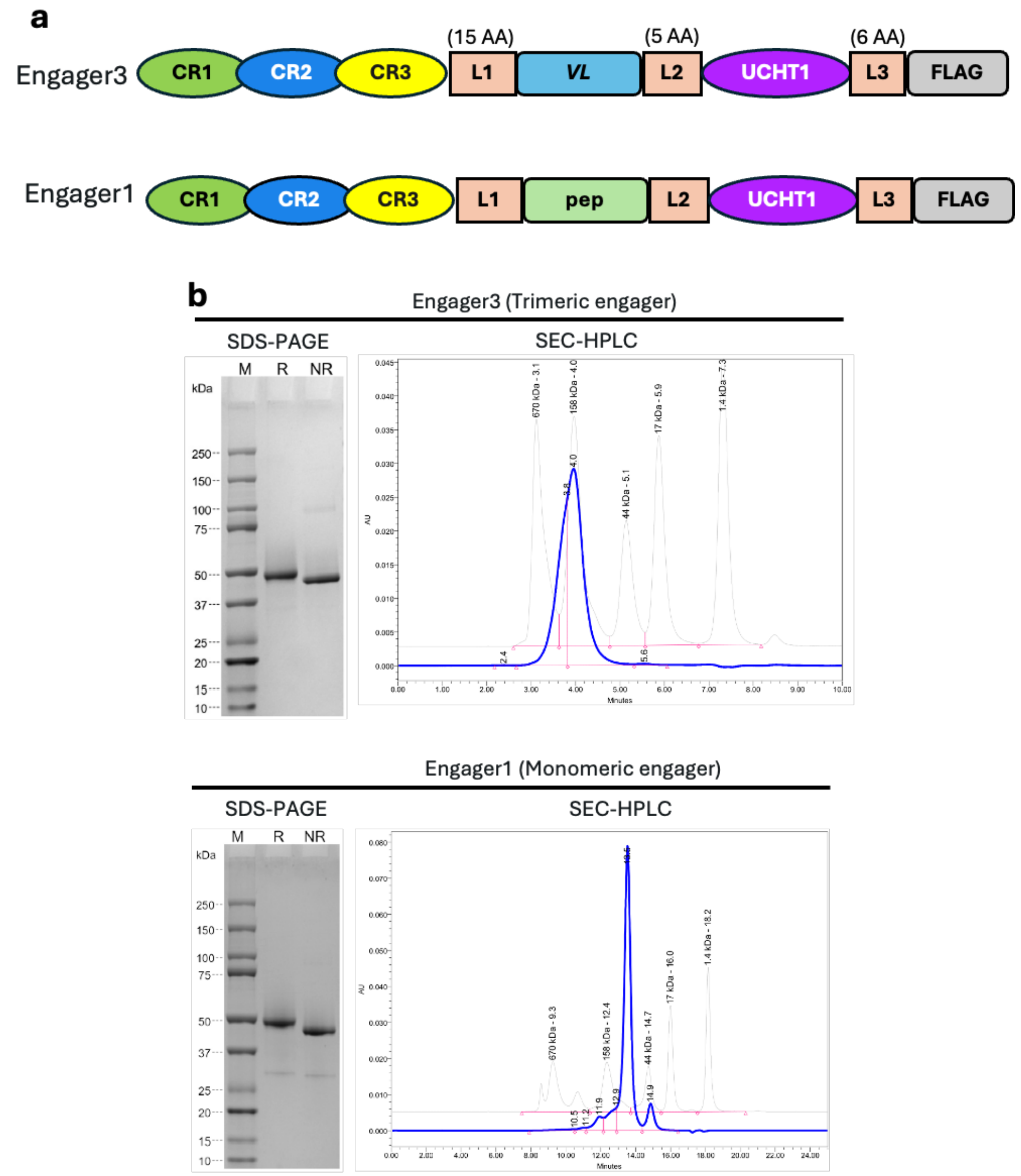


**Supplementary Fig. 4 | Design and preparation of trimeric and monomeric anti-CD3 engagers. a,** The trimeric Engager3 comprises human sLDLR$_{(25\text{-}145)}$, a $(GGGGS)_3$ linker (L1), a GCN4-derived trimerizing leucine-zipper peptide (*VL*), a GGGGS linker (L2), the CD3-targeting UCHT1 single-chain variable fragment, a GSGGSG linker (L3), and a C-terminal FLAG tag. In the monomeric Engager1 control engager, *VL* is replaced by a non-trimerizing peptide of the same length. **b,** SDS–PAGE and size-exclusion HPLC analysis of purified proteins. M, molecular-mass markers (kDa); R, reducing conditions; NR, non-reducing conditions.

### Supplementary Fig. 5

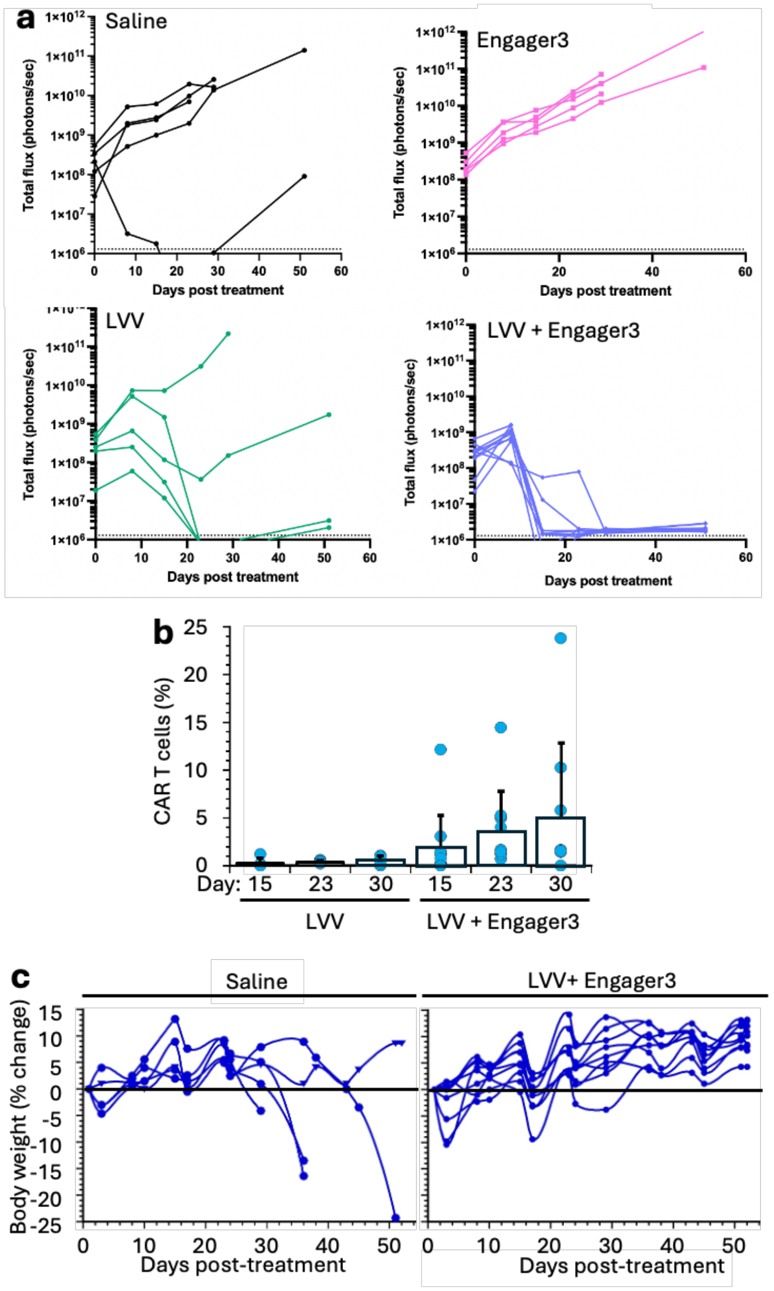


**Supplementary Fig. 5 | Circulating CAR T cells and Body weights after in vivo CAR T cell generation. a,** Bioluminescence flux (photon/sec) of OPM-2 tumors in individual mice of the four treatment groups. **b,** Percent of circulating CAR-positive T cells in the blood of mice at the indicated days post vector administration. Cells were gated on human $CD45^{+}CD3^{+}$ T cells, and quadrant values indicate the percentage of cells vs. all T cells. **c,** Body weight of tumor-bearing mice that received either PBMCs and saline, or those treated with the preformed complex of BCMA CAR-encoding LVV and trimeric engager3.
